# Human left ventricle circRNA-miRNA-mRNA network analyses reveals a novel proangiogenic role for circNPHP1 under ischemic conditions

**DOI:** 10.1101/2024.06.04.597402

**Authors:** Maryam Anwar, Moumita Sarkar, Kerrie Ford, Gianni D Angelini, Prakash Punjabi, Abas Laftah, Aránzazu Chamorro-Jorganes, Jiahui Ji, Prashant K Srivastava, Enrico Petretto, Costanza Emanueli

## Abstract

**Background:** Ischemic heart disease (IHD) is characterized by insufficient myocardial blood flow in the left ventricle and aggravated by diabetes mellitus. Endothelial resilience and reparative angiogenesis are tightly controlled processes. Gene expression is regulated by multimodal interactions between RNA species. Circular RNAs (circRNAs) can sponge microRNAs (miRNAs) to reduce the repressive effects of miRNAs on its messenger RNA (mRNAs) targets.

**Methods:** Left ventricle whole RNA-sequencing (circRNAs, mRNAs) and small RNA-sequencing (miRNAs) datasets were obtained from 3 patient groups: IHD with/out T2DM and controls (N=11 to 12/group) as part of a prospective observational cardiac surgery study.

The interactions between differentially expressed (DE) circRNAs, miRNA and mRNAs were identified with a customized bioinformatics pipeline. The emerging networks were screened using endothelial-specific RNA-sequencing datasets from GEO resulting in EC-rich networks. CircRNAs from these networks were subsequently screened (RT-PCR) in endothelial cells (ECs) exposed to disease-mimicking conditions vs control. Afterwards, circRNA pulldown allowed to interrogate the circRNA-miRNAs interactome in ECs. EC biology assays using loss-of-function and gain-of-function approaches corroborated the study.

**Results:** We identified novel circRNA-miRNA-mRNA interactions in the human diseased heart. CircNPHP1, which is upregulated in IHD with/out Type-2 diabetes mellitus (T2DM), sponges miR-221-3p to de-repress VEGF-A and BCL2, increasing the angiogenic capacity of ECs under disease and disease-mimicking conditions.

**Conclusions:** The interactions between individual members of different RNA species are affected by IHD. The therapeutic value of circNPHP1/miR-221-3p axis could be further investigated.

**Visual Abstract:** 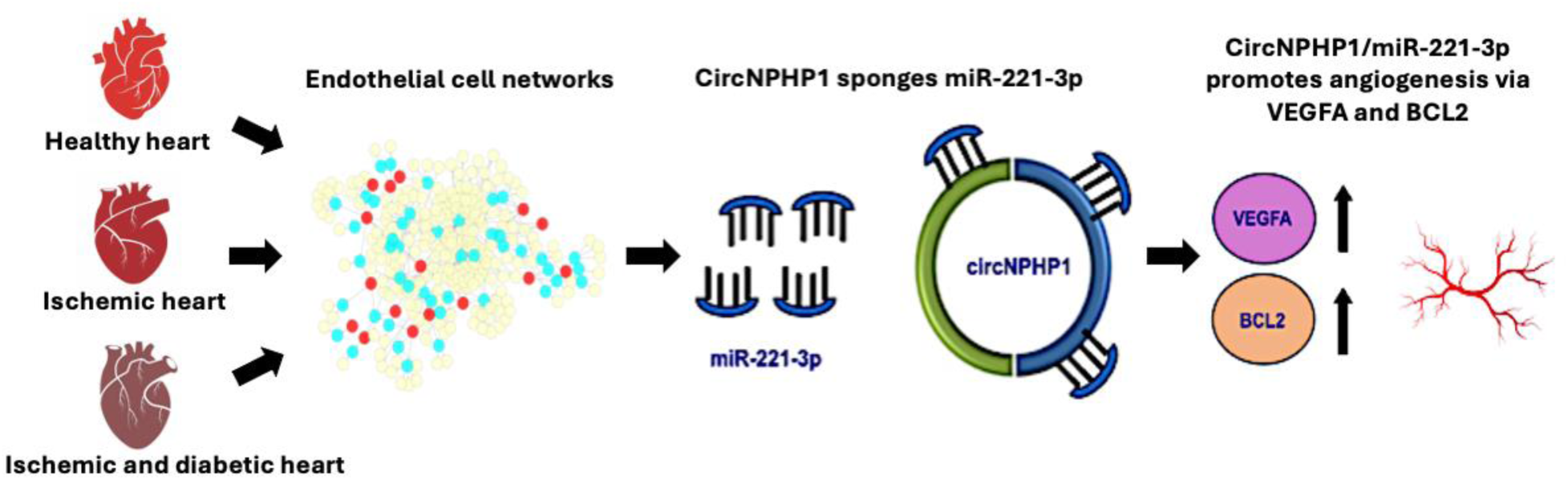

**Highlights:**

- We found a novel circRNA-miRNA-mRNA network in IHD and Type 2 diabetes.
- CircNPHP1 regulates angiogenesis and proliferation in the cardiac ECs exposed to conditions mimicking IHD and Type 2 diabetes.
- We elucidated a novel pro-angiogenic subnetwork commanded by circNPHP1/miR-221-3p/BCL2/VEGFA.
- We identified circNPHP1 as a potential new target for therapeutic angiogenesis.

## INTRODUCTION

Ischemic heart disease (IHD) can trigger adverse cardiac remodelling leading to heart failure and remains a major cause of premature death worldwide^1^. Coronary artery disease (CAD) is the prevalent cause of IHD. Diabetes aggravates CAD and impinges microangiopathy, further compromising the coronary artery and microvascular blood flow^2^. Additionally, diabetes impairs the potential for compensatory myocardial angiogenesis^3^. A prime target of hyperglycemia are the vascular endothelial cells (ECs), which line the blood vessels lumen. Endothelial dysfunction significantly contributes to pathophysiology of diabetes macrovascular and microvascular complication^4,5^. Conversely, ECs play a pivotal role in reparative angiogenesis and in maintaining the cardiac functions^6,7^. The identification and validation of novel molecular mechanisms which regulate the endothelium of the diabetic heart is of great pathophysiological significance and could additionally aid in the repurposing and design of therapeutics. Indeed, RNA therapeutics are currently in the spotlight for treatment of atherosclerosis cardiovascular disease^8^ and heart failure^9^.

CircRNAs are single stranded, covalently bonded loop structures^10^. Their circular shape makes them resistant to exonucleases and hence stabler than other RNA forms^11^. CircRNAs, which were first reported as expressed in humans in 1993, were initially considered as products of “mis-splicing”^12^. Later studies indicated their role as potential regulators of physiological functions^13,14^. CircRNAs have been implicated in post-transcriptional regulation of gene expression through the “sponging” of microRNAs (miRNAs)^15^. miRNAs are small non-coding RNAs, which can become a part of RNA-induced silencing complex (RISC) to repress messenger RNAs (mRNAs). Canonically, a miRNA recognizes its mRNA targets using its miRNA seed sequence which is semi-complementary to one or more binding sites usually placed in the 3’- UTR of the mRNAs^16,17^. CircRNAs bind to miRNAs via multiple miRNA response elements (MRE) which make the miRNAs unavailable to bind to and repress their target mRNAs^18^. This results in the de-repression of the miRNA target gene, increasing target gene’s mRNA and protein levels.

In contrast to linear RNA, circRNAs are generated through back splicing that connects the exon towards the 3’ end to one of the exons close to the 5’ end thus forming a circular structure^10^. This alteration in exon order provides a criterion for efficiently detecting back-spliced junctions during the alignment step in bioinformatics analyses^14^.

We recently reviewed the RNA regulatory role of circRNAs in IHD and cardiac remodelling^19,20^. CircRNAs have been reported to regulate cardiac angiogenesis^21^ and atherosclerosis consequent to flow-dependent endothelial inflammatory responses^21-^

^23^ in mice. Made’ et at recently investigated circRNA-miRNA-mRNA circuits in the human failing hearts using samples collected during both the surgical ventricular reconstruction procedure and heart transplantation^24^. However, the circRNAs governing endothelial-relevant miRNA-mRNA networks in the pre-heart failure left ventricle affected by IHD and T2DM remains largely unexplored. This is significant because endothelium-targeted intervention is expected to have more value at this stage of disease progression where they could be synchronized with surgical coronary revascularization.

To characterize clinically relevant circRNAs, miRNAs and mRNAs, our novel study has employed a reverse translational approach based on the use of left ventricle biopsies collected from female and male CAD patients with/out T2DM during a coronary artery bypass graft surgery (CABG) procedure. The non-ischemic, non-diabetic control group was formed by donors undergoing mitral valve repair (MVR) as sole procedure. The MVR subjects were not affected by angina, heart failure, arrhythmia or any infection and immunity conditions. Further details are provided in the Methods, when describing the ARCADIA study, which also included the pre-definition of the ARCADIA analyses plans. We have applied an integrative approach involving bioinformatics and experimental validations in the heart samples and cultured cardiac cell models.

## METHODS

### ARCADIA sample collection

ARCADIA (Association of non-coding RNAs with Coronary Artery Disease and type 2 Diabetes - REC Number: 13/LO/1687) is a prospective observational clinical study developed between two-centres, the Bristol Royal Infirmary (University Hospitals Bristol NHS Foundation Trust) and the Hammersmith Hospital (Imperial College Healthcare NHS Trust). Written informed consent was obtained from all patients. All human samples were obtained in accordance with the principles of the Declaration of Helsinki. The study was reviewed and approved by the National Research Ethics Service Committee London–Fulham (date of approval, 20 Dec 2013; reference REC 13/LO/1687), which also received our voluntary submission of the ARCADIA stage 1 analysis plan (<u>Supplementary File 1</u>). Patients’ characteristics are presented in Supplementary Table 1.

### Clinical sample collection, processing and RNA-sequencing

Left ventricular (LV) biopsies were taken from the apex of the heart using a biopsy needle. Detailed methods are available in Ford *et al*^25^, a methodological article, which reports our optimisation of the protocol for RNA extraction, rRNA depletion, libraries preparation and RNA sequencing (Illumina HiSeq 2500 and Qiagen-small RNA-seq)^25^.

### Bioinformatics analyses of RNA-sequencing data

The quality of sequenced data was assessed using the program FASTQC^26^. The library size for small RNA was estimated at 20 million reads and read length of 50-60 bp. The library size for whole transcriptome was estimated at 70 million paired-end reads with a read length of 100 bp. Samples were next aligned to human genome (hg19) using the aligner STAR^27^. Raw read counts were generated using the R package Rsubread^28,29^ and annotations were carried out using Ensembl GRCh37 (v75) (whole transcriptome) and miRBase v20 (small RNA). Raw read counts were normalised using Trimmed Mean of M-values (TMM) method, and differential expression (DE) calculated between the different groups using R package *Limma*^30^. CircRNAs were detected and annotated using CIRCexplorer^31^. CircRNAs were considered expressed in a particular group if at least 50% of the samples within that group had non-zero expression values for the circRNA. This was followed by normalisation (FPM) and DE analyses (R package *Limma*) (Supplementary Figure 1A). FC and p-values were generated using *Limma*. P-value threshold of less than 0.05 was used for finding DE circRNAs, miRNAs and mRNAs.

### Prediction of circRNA-miRNA-mRNA networks present in the human left ventricle

We predicted sponging interactions between DE circRNAs (LV biopsies) and miRNAs using three different tools/databases: TSCD (Tissue specific CircRNA database)^32^, CSCD (Cancer-Specific CircRNA Database)^33^ and CRI (Circular RNA Interactome)^34^. Next, we predicted miRNA-mRNA interactions using miRWalk suite^35^ and miRcode^36^. All interactions passing the threshold of binding probability (>0.95) were retained. Following this, we created circRNA-miRNA-mRNA networks corresponding to IHD and IHD+T2DM in Cytoscape^37^. Next, we extracted all circRNAs, miRNAs and mRNAs from the networks that were expressed in human ECs based on Gene Expression Omnibus (GEO) RNA-seq datasets: GSE100242^38^ (circRNAs, Human Umbilical Artery Endothelial Cells (HUAECs), GSE53315^39^ (miRNAs, Human Coronary Artery Endothelial Cells (HCAECs); GSE134489^40^ (mRNAs, HCAECs). Gene Ontology (GO) and KEGG (Kyoto Encylopedia of Genes and Genomes) pathway analysis was carried out using WebGestalt. Whereas GO provides information on cellular compartmentalization, biological processes and molecular function of genes, KEGG is a collection of manually drawn pathways representing information of the molecular interaction between genes and reactions for metabolic processes. KEGG pathways and GO biological processes that were significant with an FDR less than 0.05 were reported. Following pathway analysis, each of the three networks were scanned to identify circRNAs, miRNAs and mRNAs that mapped to EC-relevant pathways. From these, circRNAs that mapped to EC-relevant pathways were selected for expressional analyses in cultured ECs. The miRNA and mRNA partners of circRNAs (that showed significant differences in cultured ECs under disease-mimicking conditions) were further investigated. A pipeline indicating these steps is given in Figure 1.

**Figure 1:**
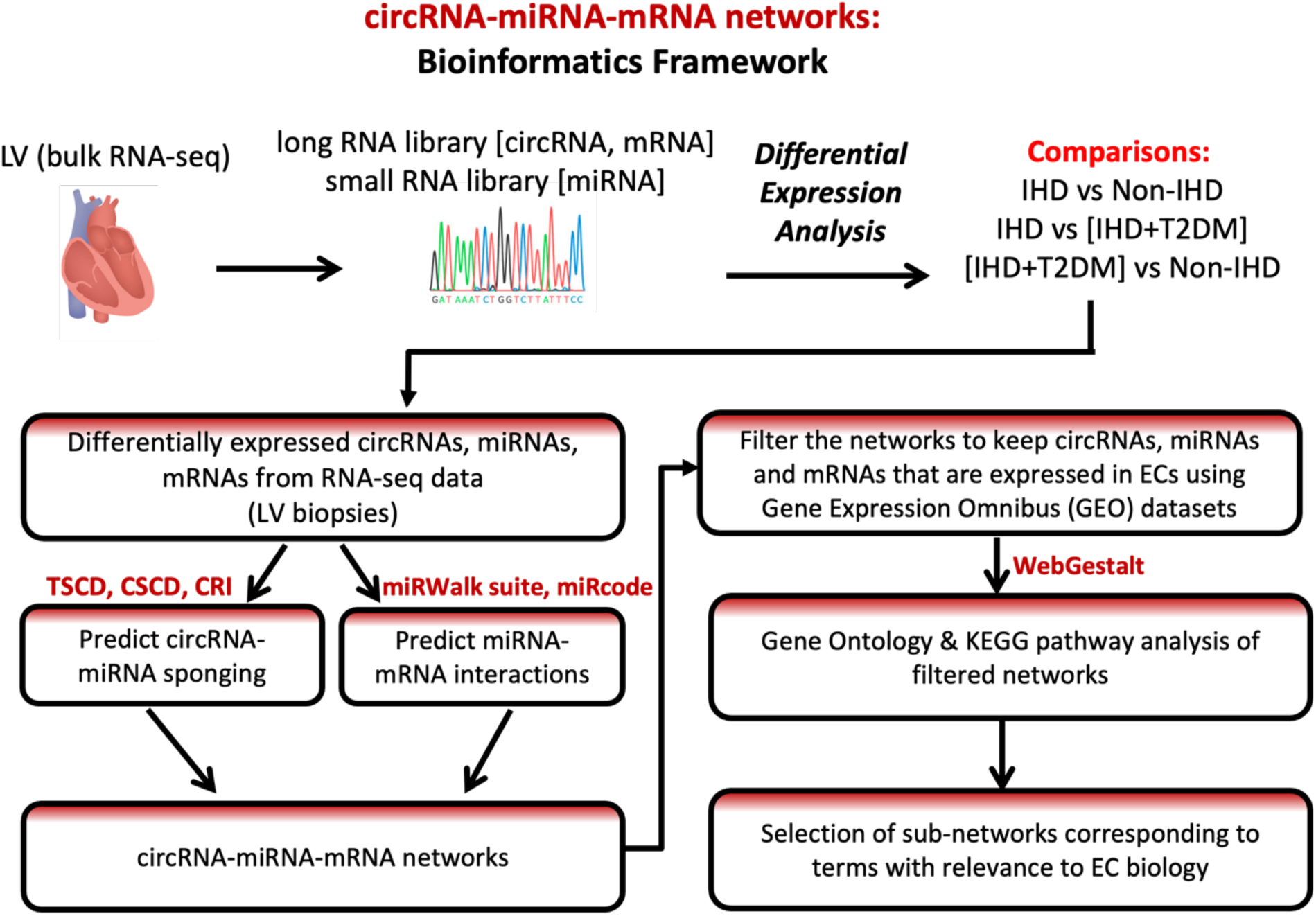
A bioinformatics framework to build circRNA-miRNA-mRNA networks. After extracting RNAs (circRNAs, miRNAs and mRNAs) from RNA sequencing data on LV biopsies of cardiac surgery patients [IHD(N=12), IHD+T2DM(N=11), non-IHD(N=12)], differential expression analyses were performed. Next, the circRNA- miRNA sponging interactions were predicted using three databases: Tissue-Specific circRNA database (TSCD; http://gb.whu.edu.cn/TSCD/), cancer-specific circRNA database (CSCD; http://gb.whu.edu.cn/CSCD) and Circular RNA Interactome (CRI; https://circinteractome.nia.nih.gov/). The miRNA-mRNA interactions were predicted using miRWalk (http://mirwalk.umm.uni-heidelberg.de/) and miRcode (http://mircode.org). Furthermore, only those interactions were kept where the connecting nodes (circRNAs, miRNA and mRNAs) were expressed in the ECs (using published EC datasets from GEO). Gene Ontology and KEGG pathway analyses were subsequently performed on these filtered networks. Sections of the network that mapped to processes corresponding to EC biology were further analysed.

### Single cell and single nuclei RNA-sequencing data analysis

The raw data for single-nuclei and single-cell RNA sequencing from healthy donor hearts was acquired from human heart atlas^41^. The pseudo counts were calculated as the sum of Unique Molecular Identifier (UMIs) in cells of each cell-type from one donor and further normalized as counts per million (CPM). Boxplots were created to show the expression across different cell types, presented as log2-transformed CPM values. We also explored single nuclei data from Koenig et al^42^, a study including 17 heart samples from people without diabetes and 9 samples from diabetic subjects. We used this data to produce boxplots of expression in diabetic vs. non-diabetic patients and performed Wilcoxon test to determine p-values for each cell-type. P-values less than 0.05 were deemed significant.

### RNA Isolation and Quantitative Real-Time Polymerase Chain Reaction (qPCR)

Total RNA was isolated from the tissue samples and cells using QIAzol reagent (Qiagen); *mir*Vana kit (Thermo Fisher Scientific) for LV biopsies and miRNeasy Mini Kit (Qiagen) for cells following manufacturer’s protocol.

For detection of circRNAs, divergent primers against the backsplice junction sequence were designed using NCBI Primer Design webtool. CircBase ID for circRNA candidates was identified according to the specific genomic coordinates. A 200nt long junction sequence, combining 100nt from the 3’ end of the circRNA sequence to 100nt from the 5’ end, was acquired using the Circular RNA Interactome webtool as the PCR amplicon sequence used for divergent primer design. For the analysis of circRNAs, cDNA was synthesized using the PrimeScript RT-PCR kit (Takara), according to the manufacturer’s instructions. Quantitative real-time polymerase chain reaction (qPCR) was performed using TB Green Premix Ex Taq Kit (Takara) on QuantStudio 6 Flex Real-Time PCR System (Life Technologies). 18S rRNA (ribosomal RNA) was used for normalization of circRNAs. For the quantification of miRNAs in the tissue samples and cells, cDNA was synthesized using miRCURY LNA RT Kit (Qiagen). Quantitative real-time polymerase chain reaction was performed using miRCURY LNA SYBR Green PCR Kit (Qiagen) following manufacturer’s instructions. Small RNA U6 was used for normalization of miRNAs. For the analysis of mRNAs, cDNA was synthesized using the PrimeScript RT-PCR kit and qPCR was performed using TB Green Premix Ex Taq Kit as well. 18S rRNA (ribosomal RNA) was used for normalization for the mRNAs.

For the quantification of circNPHP1 and linear NPHP1 in the patient plasma samples, RNA was extracted from 200 μL plasma using QIAzol and the miRNeasy Serum/Plasma Kit (Qiagen) according to the manufacturer’s instructions. 1 femtomole of external spike-in cel-miR-39 (Norgen Biotek) was added to the plasma samples during RNA extraction. As mentioned above, cDNA was synthesized using the PrimeScript RT-PCR kit and qPCR was performed using TB Green Premix Ex Taq Kit on QuantStudio 6 Flex Real-Time PCR System (Life Technologies). GAPDH was used as endogenous control and cel-miR-39 was used as spike-in control for normalization of circRNAs.

All the PCR primer sequences are listed in the resources table (Supplementary File 3).

### Cell culture

Human cardiac microvascular ECs (HCMEC) [Promocell: C-12285 (mono-doner)- Lot numbers: 440Z021.5 (male), 440Z021.4 (male), 446Z001.1 (female), 492Z009.4 (female)] and human umbilical vein ECs (HUVEC) [Promocell: C-12208 (poly-donor) - Lot numbers: 447Z004, 450Z015, 466Z022, 503Z026] were cultured on 0.2 % gelatin (Sigma-Aldrich) in endothelial cell growth medium MV and endothelial cell growth medium 2 (EGM2) (PromoCell). The biological replicates of experiments done on the ECs were carried out on cells from different lots. Human cardiac fibroblasts (HCF) [Promocell: C-12375; Lot number:491Z026.1] were cultured in fibroblast growth medium 3 (PromoCell). Proliferating AC16 human cardiomyocytes (Sigma-Aldrich SCC109, a gift from Professor Rajesh Katare, Otago University) were maintained in basal media (DMEM/F-12 with 10% FBS, 1% L-Glutamine and 1% penicillin/streptomycin). The biological replicates in AC16 and cardiac fibroblasts were carried out on cells from different passages of the same lot Cell cultures were maintained at 37 °C in a humidified atmosphere containing 5% CO_2_. For mimicking ischemic conditions, cells were exposed to hypoxia (1% O_2_ in a humidified atmosphere containing 5% CO_2_) for 48 hours at 37 °C. For mimicking diabetes hyperglycemia, cells were cultured in 25mM D-glucose (HG). Cells were cultured in hypoxia and HG to mimic the association of IHD and diabetes.

### Transfection of oligonucleotides

HCMECs and HUVECs were seeded (1×10^4^ cell per well) in a 96-well plate and transfected with 30 nM of circNPHP1 siRNA or siRNA control (non-targeting negative control pool) (Horizon Discovery). Likewise, 30 nM of hsa-miR-221-3p mimic or miRNA mimic negative control (Horizon Discovery).; anti-hsa-miR-221-3p or anti-miRNA negative control (ThermoFisher Scientific) were transfected in the ECs.

AC16 and cardiac fibroblasts were seeded (1×10^4^ cell per r well) in a 96-well plate and transfected with circNPHP1 siRNA (30 nM) or control siRNA. All the transfections were carried out using Lipofectamine^TM^ 2000 (ThermoFisher Scientific) as described^43^. Details of the sequences and catalogues are provided in the resources table <u>(Supplementary File 3)</u>. At 48 hours post transfection, cells were used for experiments described in the below subparagraphs.

### BrdU Cell Proliferation Assay and Annexin V Apoptosis Assay

After cell seeding (1×10^4^ cells well) and cell transfection, the proliferation of HCMEC, HUVEC, AC16, or cardiac fibroblasts was determined by BrdU (bromodeoxyuridine) incorporation using BrdU Cell Proliferation ELISA kit (Abcam) following manufacturer’s instructions. Cell death was measured using Annexin V Apoptosis and Necrosis Assay RealTime-Glo™ kit (Promega) in complete medium following manufacturer’s instructions.

### Endothelial Cord Formation Assay

After transfection with either circNPHP1 siRNA or control siRNA, HUVECs (13×10^4^ per well) or HCMECs (15×10^4^ per well) were seeded on a 96-well plate coated with 50 µL Growth Factor Reduced Matrigel (Corning). After 8 hours, cells were fixed with 4% formaldehyde followed by permeabilization with Triton X-100 and staining with Phalloidin (Thermo Fischer Scientific). Images were captured at ×4 magnification using a *Zeiss Axio Observer* inverted microscope. Cord formation was analyzed using the angiogenesis analyzer plugin for NIH ImageJ software.

### CircRNA pull down

The circNPHP1 pulldown was carried out in HUVECs, cardiac fibroblasts and AC16- cardiomyocytes according to the protocol described in^44^. The circRNA was pulled down using antisense oligo (ASO) probe synthesized with biotin-TEG added to the 3’ end of the sequences (Sigma-Aldrich). The first probe was designed by joining the last 15 nucleotides of circNPHP1 to the first 15 nucleotides to make a 30-nucleotide sequence against the backsplice junction followed by reverse complement. The second probe (probe 2) used to pulldown circNPHP1 consists of a 22-nucleotide sequence against the backsplice junction, (same as the siRNA sequence for circNPHP1 used in this study). Control ASO was chosen following the protocol The sequences of the probes are listed in the resources table <u>(Supplementary File 3)</u> and additionally indicated in Supplementary Figure 4A.

The brief protocol for pulling down the circNPHP1 was the following. For each cell type, 5 million cells were harvested in Tris, KCl, MgCl_2,_ nonidet P-40-polysome extraction buffer (PEB). Cell extracts were incubated with 1µl of the biotinylated probes or the control probe. (100 µM) in Tris, EDTA, NaCl, Triton (TENT) buffer with rotation for 90 minutes at 4°C for hybridization. Subsequently, streptavidin beads (washed in TENT buffer) were added and the mixture of the cell extract, probes and the beads were incubated with rotation for further 45 minutes at room temperature. Then the beads were washed thrice in TENT buffer followed by isolation of RNA using Trizol and phenol-chloroform. cDNA was synthesized and subjected to qPCR for determination of circNPHP1 and the bound miRNAs. Fold enrichment was calculated in the pulldown samples with probe 1 and probe 2 compared to the control probe using delta delta Ct method (Livak). 18S was used as reference gene to normalize for circNPHP1 and linear NPHP1 and U6 was used as to normalize for the miRNAs. In HUVECS, four miRNAs were screened namely miR-221-3p, miR-222-3p, miR-299-3p and miR-141-3p. Subsequently, in the pulldown assays performed in cardiac fibroblasts and AC16, miR-221-3p was checked for binding to circNPHP1.

### Western blot analyses

Cells were harvested and lysed in ice-cold RIPA buffer (Sigma-Aldrich) containing 1 mM orthovanadate, 1 mg/mL of protease inhibitor cocktail (Roche). Protein concentrations were determined using Bradford assay reagent (BIORAD). Equal amounts (15μg) of proteins were fractionated in 4-20 % SDS-polyacrylamide gel electrophoresis (SDS-PAGE) and transferred onto a polyvinylidene difluoride (PVDF) membrane using wet transfer method. The membranes were blocked with 5% milk followed by incubation with primary antibodies overnight using the following antibodies: anti-VEGF-A (Abcam), anti-BCL2 (Cell Signaling Technology), anti-Lamin B1 (Cell Signaling Technology). The membrane was further probed with anti-Rabbit IgG-HRP (Santa Cruz Biotechnology). Immobilon Crescendo Western HRP substrate (Merck Millipore) was used to detect the chemiluminescence. Protein bands were visualized using the ChemiDoc^TM^ MP Imaging System (BIORAD). Densitometry analysis of the gels was carried out using NIH ImageJ software. The details of antibodies are listed in the resources table.

### Statistical analyses and data management

The analyses of the RNA-seq datasets are included in the “Bioinformatics analyses of RNA-sequencing data” and “Single cell and single nuclei RNA-sequencing data analysis” sections of this file.

For the remaining data, statistical analyses were performed with GraphPad Prism software (GraphPad). Normal distribution of each experimental group was determined by Shapiro-Wilk test. Statistical significance between two groups was analysed by unpaired Student *t* test. For the comparison between multiple groups, 1-way ANOVA was performed and adjusted for multiple comparisons using post hoc Dunnett’s test. Data are presented as mean±SEM. P values ≤ 0.05 were considered statistically significant.

In adherence to FAIR policy of data management^45^, the data created and/or analysed in the current study will be made available through online repositories (see data availability section).

## RESULTS

### Differentially expressed circRNAs, miRNAs and mRNAs in LV biopsy samples

From RNA-seq analyses, the number of detected circRNAs as well as common and uniquely expressed circRNAs among the three patient groups are shown <u>in</u> Supplementary Figure 1 B. We next performed differential expression analyses and identified the significantly DE circRNAs (Figure 2A, Supplementary Figure 1C), miRNAs (Figure 2B, Supplementary Figure 1C) and mRNAs (Figure 2C, Supplementary Figure 1C) (P-value<0.05) in IHD vs non-IHD, IHD+T2DM vs non-IHD and IHD+T2DM vs IHD. The files containing lists of significant DE circRNAs, miRNAs and mRNAs relevant to each comparison between patient groups are provided as <u>Supplementary files 2a, 2b and 2c</u>).

**Figure 2:**
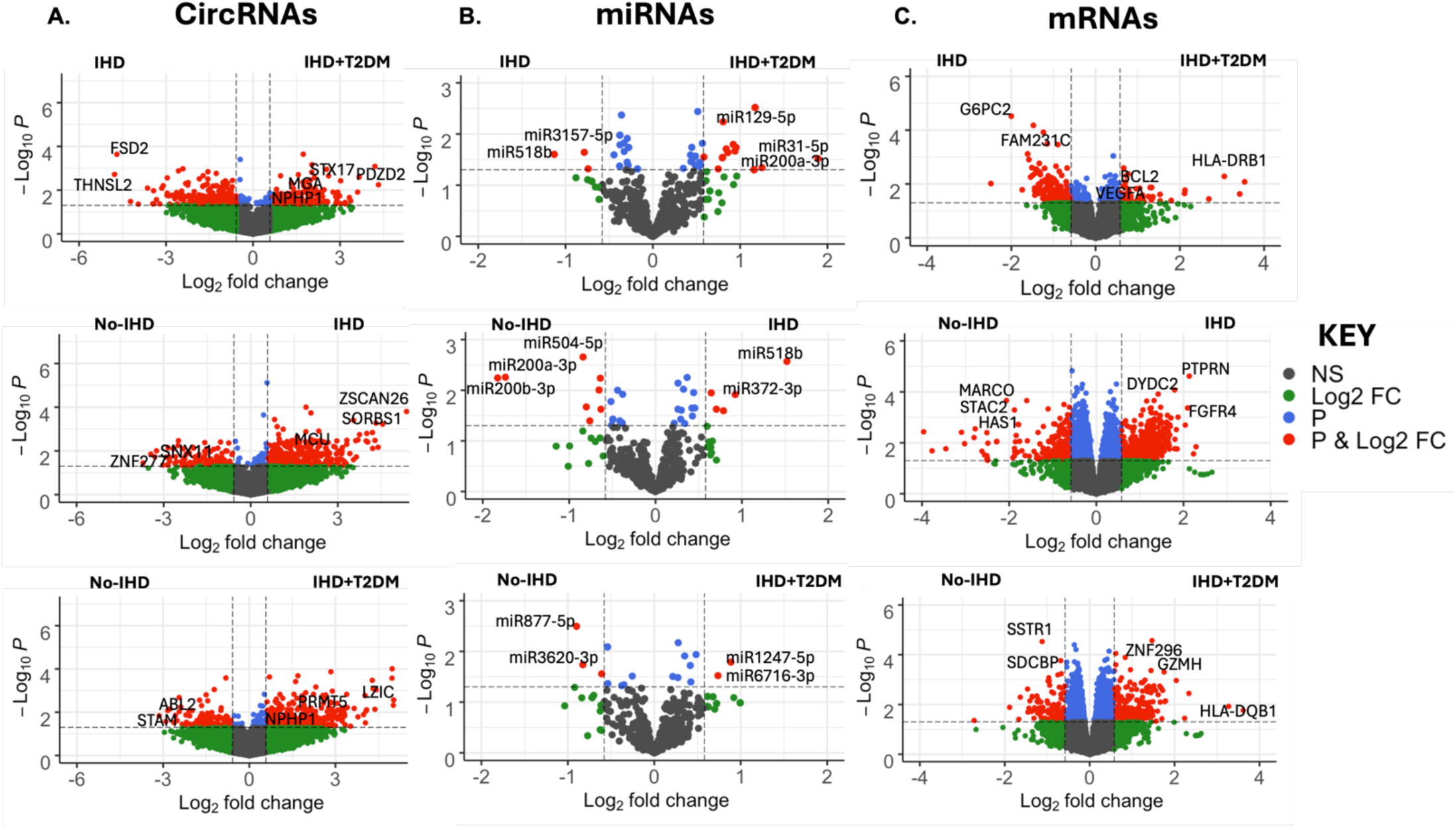
Differential expression analyses. (A-C) CircRNAs, miRNAs and mRNAs were identified following the pipeline outlined in methods [IHD(N=12), IHD+T2DM(N=11), non-IHD(N=12)]. Differential expression analyses were performed on normalised and log-transformed counts using the R package *Limma.* Three comparisons were performed (IHD+T2DM vs IHD), (IHD vs non-IHD) and (IHD+T2DM vs non-IHD). The corresponding differentially expressed circRNAs, miRNAs and mRNAs were plotted in a volcano plot with log2 fold change (FC) on the x-axis and-log10 Pvalues on the y axis. CircRNAs, miRNAs and mRNAs that passed the log2 FC threshold of +/-0.58 and P-value < 0.05 are shown in red, those that just pass the FC threshold are in green and those that only pass the P-value threshold are in blue.

### CircRNAs are predicted to sponge miRNAs that affect mRNAs involved in EC- relevant pathways

Figure 3 shows the Cytoscape representations of the interactions (“edges”, shown as connecting lines) between the circRNAs, miRNAs and mRNAs (shown as points called “nodes”). The networks indicate putative regulatory relationships between circRNAs, miRNAs and mRNAs for the three different comparisons between patient groups. Additionally, the three networks that are presented in Figure 3 have also been filtered based on expression of the RNAs in ECs by using published data on ECs from GEO (see methods for details). Figure 3 thus illustrates the following: IHD vs non-IHD (EC-enriched, 305 nodes and 359 edges - Figure 3A<u>, left</u>), IHD + T2DM vs non-IHD (EC-enriched, 1,046 nodes and 2,803 edges - Figure 3B<u>, left</u>) and IHD + T2DM vs IHD (EC-enriched, 303 nodes and 361 edges - Figure 3C<u>, left</u>). Following the networks creation, we looked at significant biological processes and pathways that these RNAs are expected to contribute to. All processes and pathways that were enriched in the network with FDR<0.05 were considered significant (Figure 3 A, B and C; right). Significant processes and pathways include VEGF signaling, angiogenesis, apoptosis and HIF-1 signaling which are of possible relevance for vascular biology and disease (highlighted in colors in Figure 3).

**Figure 3:**
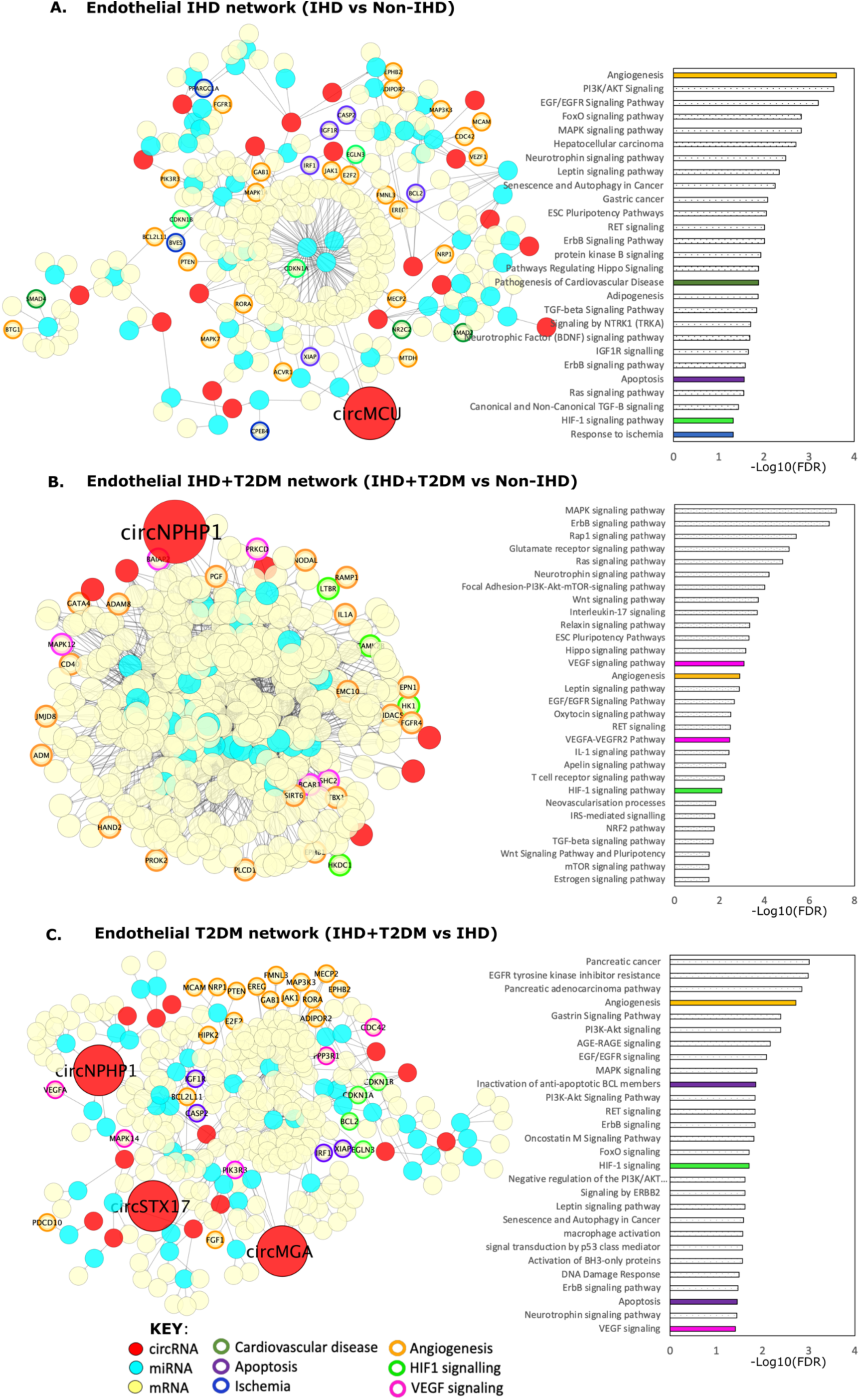
Predicted circRNA-miRNA-mRNA networks in the heart ECs. (A-C) The endothelial circRNA-miRNA-mRNA networks were constructed in Cytoscape using predicted interactions between circRNAs, miRNAs and mRNAs in IHD (+/- T2DM) (using in-house LV biopsies and heart ECs [GEO data]). All nodes (circRNAs, miRNA and mRNAs) shown in the network are expressed in LV biopsies and ECs. The networks show circRNAs in red, miRNAs in cyan and mRNAs in yellow. Following Gene Ontology (GO) and KEGG pathway analyses, all significant GO biological processes and KEGG pathways (passing threshold of FDR ˂ 0.05) were plotted as a bar graph with-log10(FDR) on the x-axis. Moreover, pathways relevant to EC biology are highlighted in the graph as solid colors. The corresponding genes (downstream targets of circRNAs and miRNAs) are outlined with the color of the relevant pathways. Genes that correspond to more than 1 pathway are outlined with multiple colors. The top significantly upregulated circRNAs that map to EC-relevant pathways are enlarged and labelled. (A) Network constructed from IHD vs non-IHD comparison. CircMCU emerged as the top significantly upregulated circRNA which together with its miRNA and mRNA partners mapped to EC-relevant pathways including apoptosis, angiogenesis and HIF1 signalling. (B) Network constructed from IHD+T2DM vs non-IHD comparison. CircNPHP1 emerged as the top significantly upregulated circRNA which together with its miRNA and mRNA partners mapped to EC-relevant pathways including VEGF signalling, angiogenesis and HIF1 signalling. (C) Network constructed from IHD+T2DM vs IHD comparison. CircNPHP1, circSTX17 and circMGA emerged as the top significantly upregulated circRNAs which together with their miRNA and mRNA partners mapped to EC-relevant pathways including VEGF signalling, apoptosis, angiogenesis and HIF1 signalling.

### Screening and validation of circRNAs predicted to be commanding EC networks involved in EC survival regulation

From each network, we selected circRNAs involved in EC-relevant enriched pathways such as angiogenesis and VEGF signalling (large red labelled nodes in Figure 3 A, B and C, left). These included circNPHP1 (Figure 3B and 3C), circMCU (Figure 3A), circSTX17 and circMGA (Figure 3C). Following pilot expressional analyses in cultured HCMECs, circNPHP1 was confirmed to increase in response to hypoxia with/out HG vs control (Supplementary Figure 1D). This resembles the comparisons of Figure 3B and the results of bioinformatics analyses hence circNPHP1 was selected for continuing the study. The regulation of circNPHP1 expression by either hypoxia or hypoxia+HG (vs standard culture conditions) was then confirmed using different batches of HCMECs and HUVECs (Figure 4C). By contrast, the linear NPHP1 form was not affected by difference in culture conditions (Supplementary Figure 3A). The data obtained in ECs exposed to disease-mimicking conditions were aligned with the results of qRT-PCR analyses in ventricle samples from the ARCADIA patients (Figure 4B), which were used to validate the circNPHP1 and linear NPHP1 results of the initial LV RNA-seq analyses (Figure 4A and 4B). The PCR analysis did not confirm the RNA-seq change for circNPHP1 between IHD with/out T2DM and identified circNPHP1 to be increased in IHD with/out T2DM vs non-IHD control.

**Figure 4.**
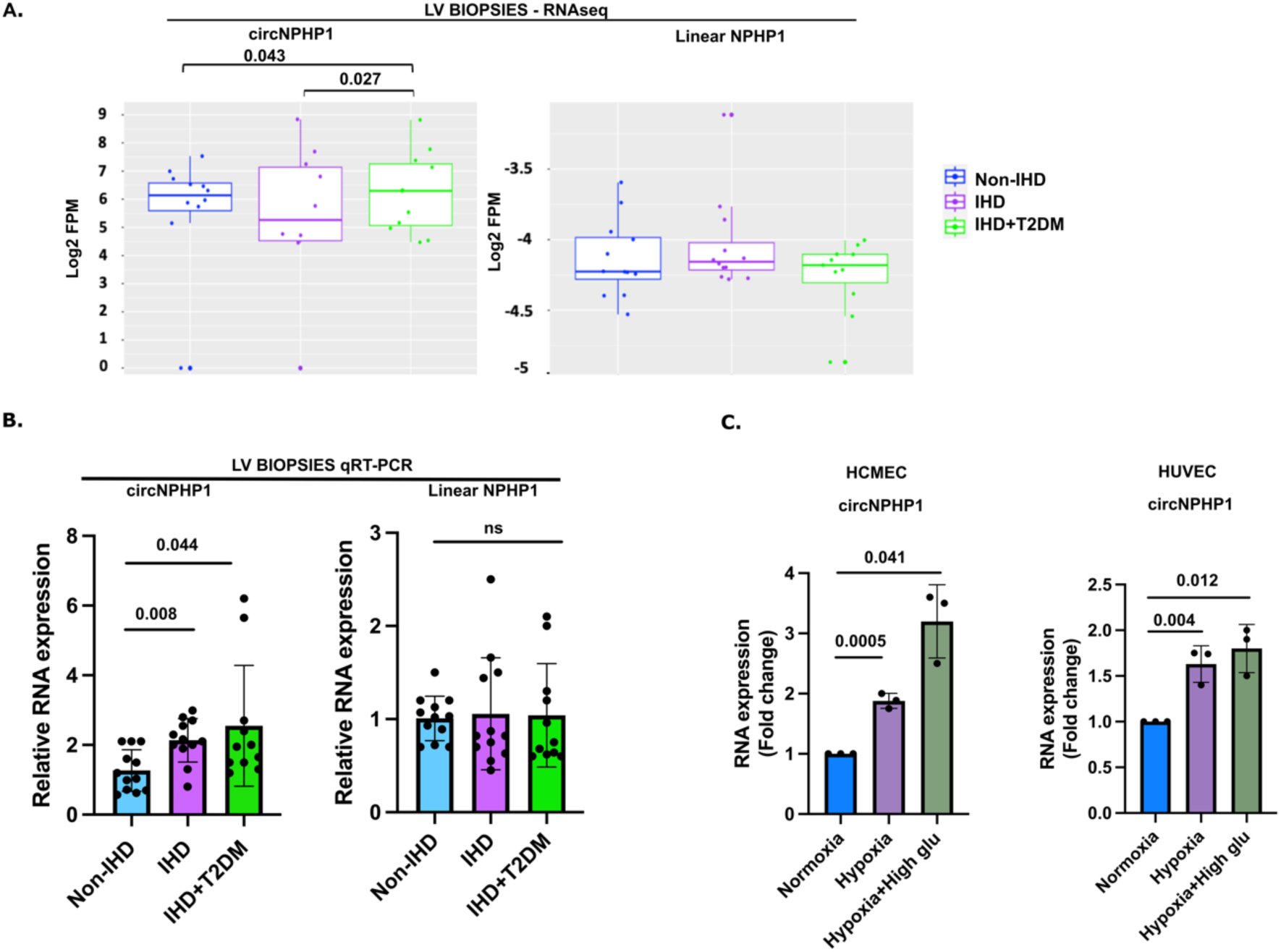
CircNPHP1 expression in the human left ventricle and cultured endothelial cells. Cardiac CircNPHP1 levels are higher in patients suffering from IHD with/out associated T2DM and in cultured endothelial cells (human cardiac microvascular endothelial cells-HCMECs-and human umbilical vein endothelial cells-HUVECs) exposed to disease-mimicking conditions, such as hypoxia and hypoxia combined with increased D-glucose level (High glu). (A) RNAseq: Log-transformed normalised (FPM) values of circNPHP1 and linear NPHP1 are plotted as boxplot. CircNPHP1 is significantly upregulated in IHD+T2DM as compared to IHD and non-IHD (P-value˂0.05). Linear NPHP1 is expressed at very low levels and does not change between the different patient groups [IHD(N=12), IHD+T2DM(N=11), non-IHD(N=12)]. (B) Quantitative real-time polymerase chain reaction (qRT-PCR) analysis of circNPHP1 (left panel) and linear NPHP1 (right panel) of LV biopsies from non-IHD (N=12), IHD (N=12), IHD+T2DM (N=11). 18S is used as housekeeping gene. Results were assessed by Kruskal-Wallis test followed by Dunn’s multiple comparison to non-IHD group. P values are indicated accordingly, P>0.05 is indicated as ns [nonsignificant]. (C) HCMEC (left panel) and HUVEC cells (right panel) were cultured in either normal conditions, hypoxia (1% Oxygen) or hypoxia combined with high glucose (25Mm D-glucose). After 48 hours, cells were harvested for qRT-PCR for the analysis of circNPHP1 (N=3). Fold change in RNA expression is relative to normoxia; 18S is used as housekeeping gene. Results were assessed by 1-way ANOVA (Dunnett’s post hoc test). P-values are indicated accordingly, P>0.05 is indicated as ns [nonsignificant].

We next used single-cell and single-nuclei RNAseq data from heart atlas (Supplementary Figure 2A) to investigate whether NPHP1 expression is confined to EC or present in different cell types of the human heart. Because traditional scRNA-seq does not allow for circRNA identification, we resolved to search for the linear form under the assumption that cells expressing linear NPHP1 would be capable to produce its circular counterparts. Our analyses identified NPHP1 to be ubiquitously expressed in the several cardiac cell types (Supplementary Figure 2B). Using single nuclei data from Koenig et al^42^, we found that expression levels of linear NPHP1 do not change between diabetic and healthy controls (P-value>0.05 for all cell types; Supplementary Figure 2C).

### CircNPHP1 promotes proliferation and capillary-like cord formation capacity of cultured ECs

The functional relevance of the endogenous circNPHP1 for the human endothelium was confirmed in HUVECs and HCMECs transfected with circNPHP1 siRNA or control siRNA and submitted to EC biology tests performed under normal, hypoxia and hypoxia+HG conditions. Knockdown of circNPHP1 specifically downregulated the circular NPHP1 form whereas linear NPHP1 remained unchanged, validating the specificity of the siRNA for the circular form of NPHP1 (Supplementary Figure 3A).

The knock-down (KD) of circNPHP1 was associated with a marked reduction of the cell proliferation as measured by BrdU incorporation across all the conditions (Figure 5A). Upon circNPHP1 knockdown in the HUVECs, cord formation assay on Matrigel showed prominent reduction of angiogenesis under standard culture conditions (Figure 5C) and as well as in hypoxia and hypoxia+HG conditions. The total tube length and the number of branches, meshes, nodes, junctions exhibited significant reduction in the HUVECs with circNPHP1 knockdown (Figure 5C and Supplementary Figure 3B). Similar effect was observed in HCMECs (<u>Supplementary 3C</u>).

**Figure 5.**
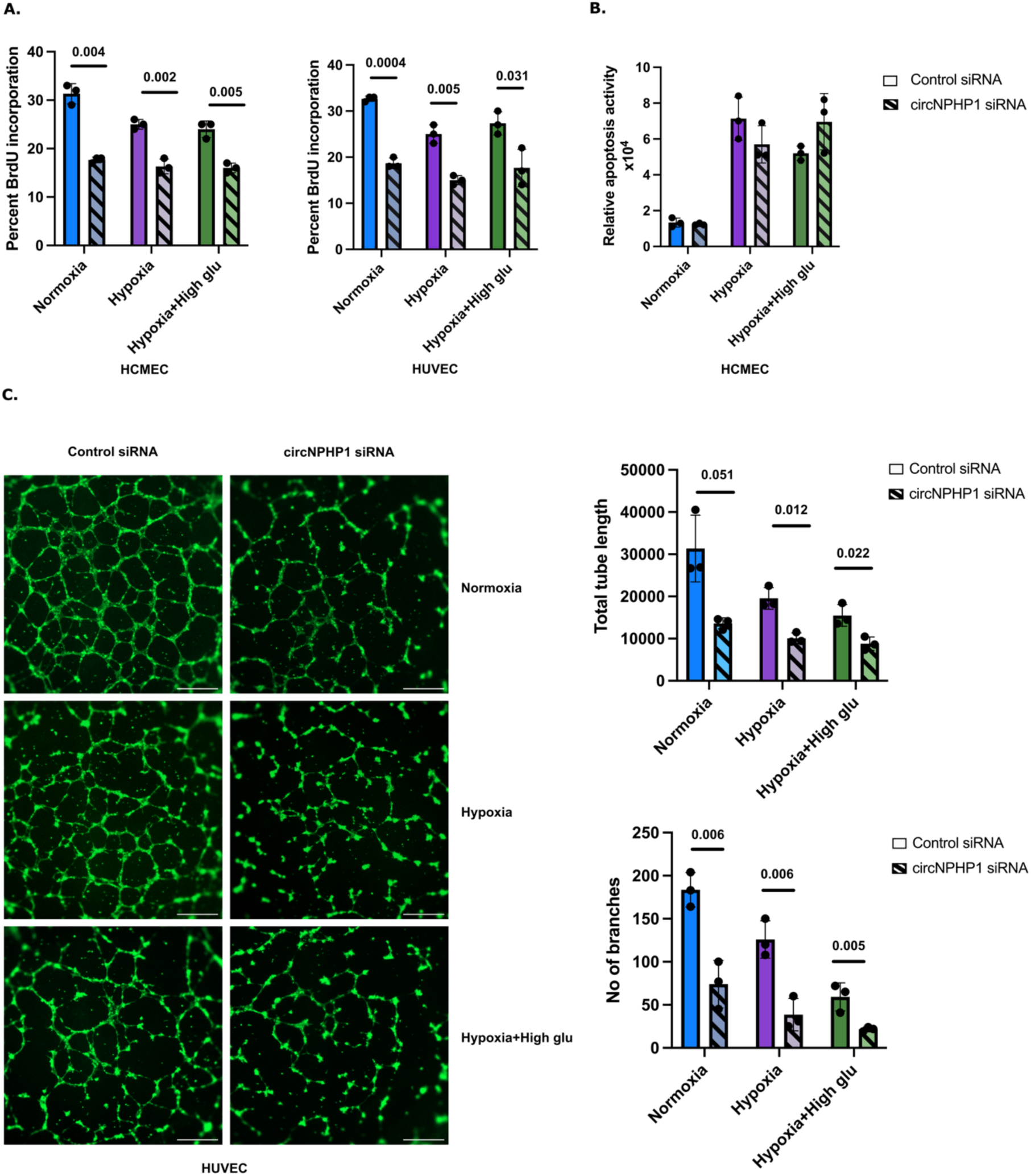
Impact of endogenous CircNPHP1 on endothelial cell proliferation, apoptosis and cord formation. Endogenous CircNPHP1 promotes the proliferative and networking capacities of HCMECs and HUVECs. (A) HCMECs (left panel) and HUVECs (right panel) were transfected with 30nM of circNPHP1 siRNA or control siRNA, and cultured in normal conditions, hypoxia (1% Oxygen) and hypoxia-high glucose (25mM D-glucose) conditions respectively. At 48 hours post transfection, cells were harvested for proliferation assay (BrDU incorporation) (N=3), and (B) HCMEC: apoptosis assay (Annexin V assay) (N=3). (C) HUVECs were transfected with 30nM of circNPHP1 siRNA or control siRNA, and cultured in either normal conditions, hypoxia (1% Oxygen) or hypoxia combined with high-glucose (25mM D-glucose) conditions. At 48 hours post transfection, the cells were seeded on a 96-well plate containing Growth Factor Reduced Matrigel and cultured in similar conditions for 8 hours to determine their capacity to network in a cord formation assay, which is a classical angiogenesis assay. Representative images of the cord formation (stained with Phalloidin, scale bar: 500 μM) (left panel). Histograms depicting the total tube length and number of branches quantified from the angiogenesis assay (right panel) (N=3). Results in (A–C) were by unpaired Student *t* test between control siRNA and circNPHP1 siRNA among the three groups respectively. P-values are indicated accordingly, P>0.05 is indicated as ns [nonsignificant].

Additionally, in hypoxia and hypoxia+HG conditions, similar reduction of angiogenesis was observed in HUVECs upon silencing circNPHP1 (Figure 5C). By contrast, apoptotic death of the cells remained unaffected upon circNPHP1 knockdown in HCMECs (Figure 5B). Altogether our results contribute to support the hypothesis that circNPHP1 is involved in promoting EC proliferation and angiogenesis.

### CircNPHP1 binds to and sponges miR-221-3p in ECs

CircNPHP1 as well as its partner miRNAs and mRNAs in the T2DM network mapped to a number of EC-relevant pathways including angiogenesis, VEGF signalling and HIF1 signalling which indicates that circNPHP1 might be involved in various pathways through a combination of miRNA and mRNA partners. We, therefore, extracted the sub-network of circNPHP1 from T2DM network. 9 miRNAs were predicted to be sponged by circNPHP1 and 38 mRNAs to be potential target genes of the 9 miRNAs (Figure 6A<u>, left</u>).

**Figure 6.**
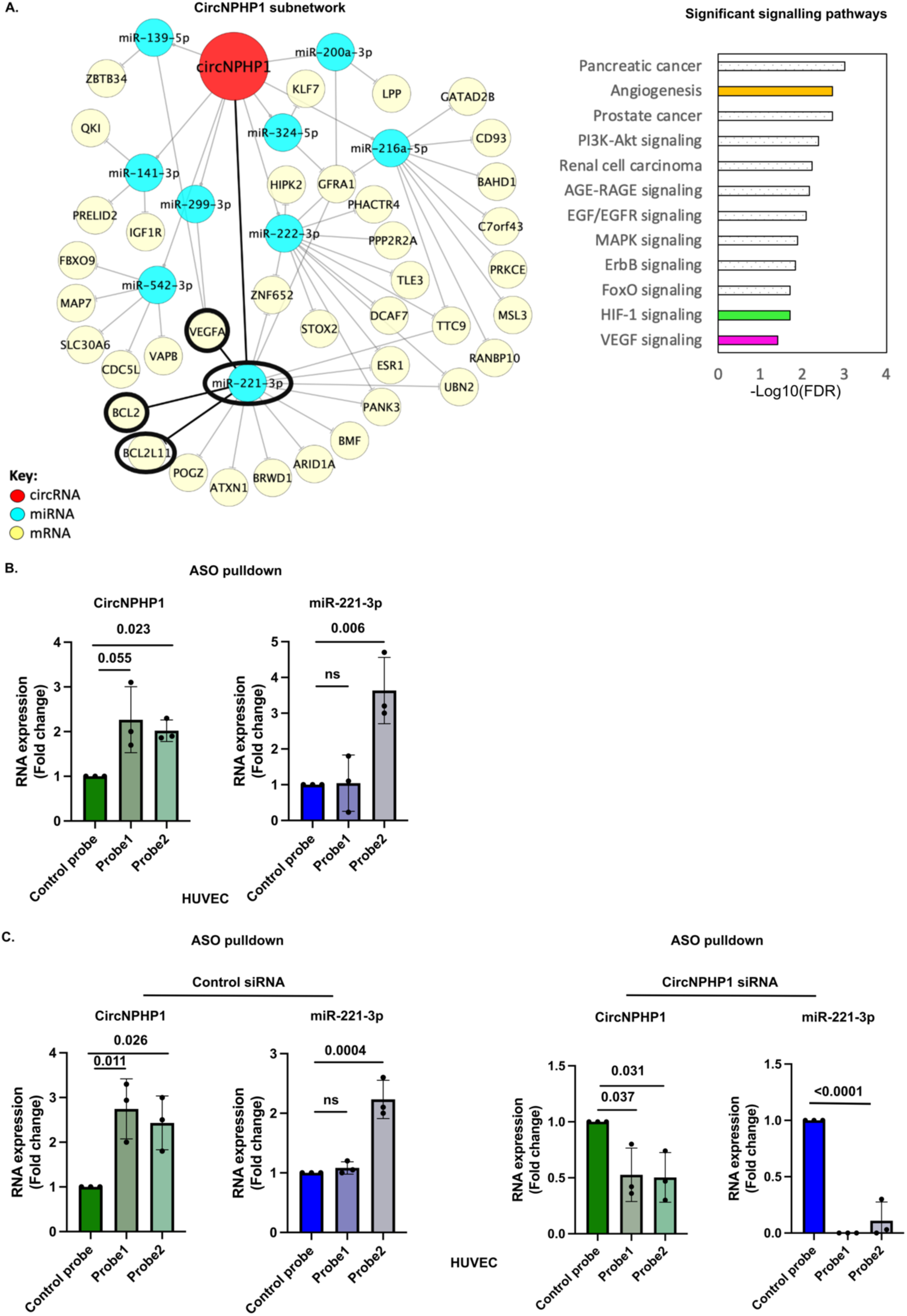
Identification and validation of the CircNPHP1 proangiogenic interactome in endothelial cells. CircNPHP1 is physically associated with the anti-angiogenic miR-221-3p to putatively command a pro-angiogenic sub-network. (A) CircNPHP1 was found to be enriched in endothelial IHD+T2DM as well as T2DM networks. In particular, circNPHP1 in T2DM network was connected to many miRNAs and mRNAs that mapped to EC-relevant pathways (VEGF signalling and angiogenesis). We, therefore, extracted the miRNAs and mRNAs that were connected to circNPHP1 from T2DM network to analyse the interactions in greater detail. The arc consisting of circNPHP1, its sponged miRNA partner (miR-221-3p) and downstream targets that were validated (VEGFA, BCL2, BCL2L11) are bold in black. (B) HUVEC cell extracts were incubated with biotinylated ASO control probe, probe 1 and probe 2 respectively (see details in methods) for pulldown of circNPHP1 using streptavidin beads. Subsequently, RNA was isolated and subjected to qRT-PCR for the analysis of circNPHP1 (left panel) and miR-221-3p (right panel). Fold enrichment was calculated in the pulldown samples with probe 1 and probe 2 against the control probe pulldown. 18S and U6 were used as reference genes to normalize for circNPHP1 and miR-221-3p respectively (N=3). (C) HUVECs were transfected with 30nM of circNPHP1 short interfering RNA (siRNA) or control siRNA. 48 hours post transfection, cells were harvested and incubated with biotinylated ASO control probe, probe 1 and probe 2 respectively for pulldown of circNPHP1 using streptavidin beads. Subsequently, RNA was isolated and subjected to qRT-PCR for the analysis of circNPHP1, linear NPHP1 and miR-221-3p. Fold enrichment was calculated in the pulldown samples with probe 1 and probe 2 against the control probe pulldown. 18S and U6 were used as reference genes to normalize for circNPHP1, linear NPHP1 and miR-221-3p respectively (N=3). Results were assessed by 1-way ANOVA (Dunnett’s post hoc test). P-values are indicated accordingly, P>0.05 is indicated as ns [nonsignificant].

After pull-down of circNPHP1 in HUVECs, we confirmed enrichment of the circRNAs (Figure 6B) while the linear NPHP1 was only detectable in the whole cell extracts and was undetectable in the corresponding circNPHP1 enriched conditions, indicating specific pulldown of the circular RNA (Supplementary Figure 4C). After pull-down, out of the 9 predicted miRNAs, only miR-221-3p appeared to be enriched and hence bound with the circRNA (Figure 6B and Supplementary Figure 4B). Of note, the miR- 221-3p enrichment was effective in the case of probe-2 but not in probe 1. A possible explanation could be steric hindrance in binding of miR-221-3p caused by other factors bound to probe1 due to larger size. Importantly, knocking down the circNPHP1 in HUVCs before pulling-down for the circRNA abolished the enrichment of miR-221-3p, strengthening the evidence of binding between miR-221-3p and circNPHP1 (Figure 6C).

To investigate if the putative circNPHP1/miR-221-axis was relevant to other cardiac cell types, we first measured the expression of both RNA molecules by qRT-PCR in human AC16-cardiomyocytes and human cardiac fibroblasts, before proceeding with the circNPHP1 pull-down experiment. Expression of circNPHP1 and miR-221-3p in AC16 was well-detected and comparable to HUVECs (Mean ct: 30 for circNPHP1; Mean ct: 19 for mir-221-3p). Although miR-221-3p exhibited good expression (Mean ct: 25) in the cardiac fibroblasts, the expression of circNPHP1 as well as the linear form was poor (Mean ct: 35) (Supplementary Figure 4D). In AC16, probe 2 induced effective pulldown of circNPHP1, which was associated with enrichment of miR-221- 3p (Supplementary Figure 4E). Due to poor expression of circNPHP1 in cardiac fibroblasts, the pulldown of circNPHP1 by either probe was undetectable. Despite the ineffective pulldown of circNPHP1, we still found enrichment of miR-221-3p in cardiac fibroblasts by both the probes (Supplementary Figure 4F). This could be possibly due to significant binding of circNPHP1 to miR-221-3p which caused enrichment in miR- 221-3p even with slight pulldown of circNPHP1. Interestingly, the proliferation of cardiac fibroblasts significantly decreased upon silencing circNPHP1 (Supplementary Figure 5A) whereas the cellular apoptosis in cardiac fibroblasts and AC16 remained unchanged (Supplementary Figure 5B). Together, our data suggests that the circNPHP1/ miR-221-3p axis is operational in different cell types and could mediate different responses, which will need to be addressed in future studies.

### CircNPHP1 levels influence the expression of miR-221-3p target genes BCL2 and VEGFA in ECs

To confirm that sponging of miR221-3p by circNPHP1 prevents the miRNA from repressing target mRNAs, we focussed on three targets of miR221-3p emerging from literature: BCL2, BCL2L11 (BIM) and VEGFA^46,47,48,49^. These 3 genes were already included in the angiogenesis and cell survival pathways emerging from our pathway analyses in heart biopsies (FDR < 0.05). We validated that miR-221-3p mimic repressed VEGFA and BCL2 expression in HUVECs, while anti-miR-221-3p produced the expected opposite effects. By contrast, BIM was unaffected by the forced changes in miR-221-3p expression (Supplementary Figure 6A and 6B). Independently on the culture conditions, in HUVECs with circNPHP1 KD, the VEGFA and BCL2 mRNA levels decreased (vs HUVECs transfected with the siRNA-control) (Figure 7A), while BIM remained unchanged (Figure 7A). Therefore, BIM was excluded from further analyses. Western blots for VEGFA and BCL2 showed that protein expression was aligned with the mRNA expressional data (Figure 7B). To investigate if the changes in VEGFA and BCL2 expression induced by circNPHP1silencing are mediated by the release of miR221-3p repression, HUVECs were transfected with circNPHP1 siRNA or control siRNA in combination with an anti-miR221-3p. The inhibitory effect of circNPHP1 silencing on BCL2 and VEGFA was reduced when ECs were treated with anti-miR221-3p further confirming the joint operation of circNPHP1 and miR-221-3p in regulating VEGFA and BCL2 (Figure 7C). Silencing circNPHP1 in the presence of a miR221-3p mimic decreased VEGFA and BCL2 mRNA levels, without showing a further reduction of mRNA expression in comparisons to circRNA KD alone (Figure 7D).

**Figure 7.**
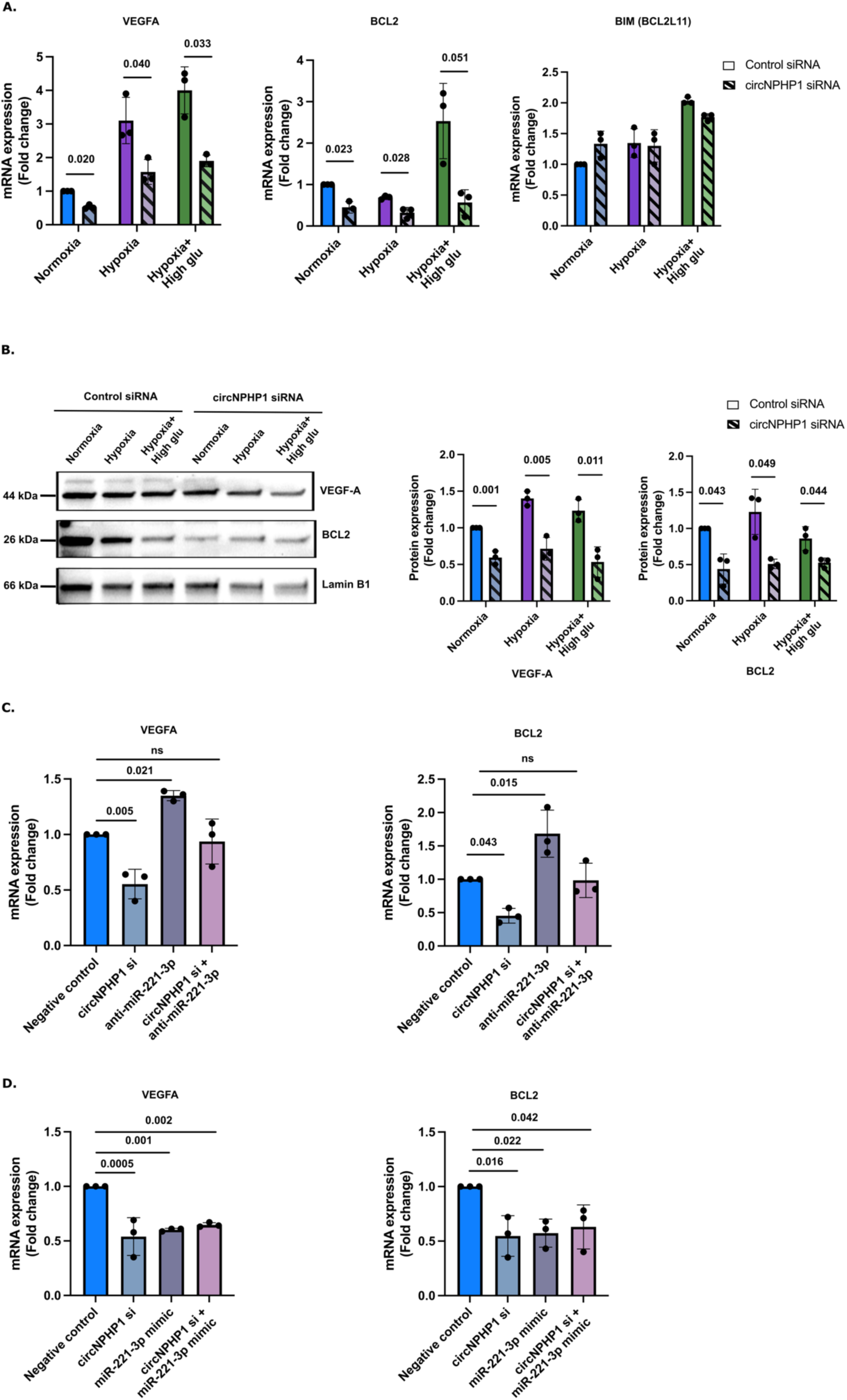
CircNPHP1 regulation of miR-221-3p target genes’ expression. (A) HUVECs were transfected with 30nM of circNPHP1 siRNA or control siRNA, and cultured in normal conditions, hypoxia (1% Oxygen) and hypoxia-high glucose (25mM D-glucose) conditions respectively. 48 hours post transfection, cells were harvested for qRT-PCR for the analysis of VEGFA, BCL2 and BIM (N=3). Fold change in mRNA expression is relative to normoxia; 18S is used as housekeeping gene. (B) HUVECs were transfected with 30nM of circNPHP1 siRNA or control siRNA, and cultured in normal conditions, hypoxia (1% Oxygen) and hypoxia-high glucose (25mM D-glucose) conditions respectively. 48 hours post transfection, cells were harvested for western blot analysis of VEGF-A and BCL2 (N=3). Images shown are representative (left panel). Histograms showing western blot quantification is expressed as fold change against normoxia and normalized to Lamin B1 (loading control) (right panel). (C) HUVECs were transfected with 30nM of circNPHP1 siRNA, anti-miR-221-3p respectively or in combination. Combination of control siRNA and anti-miRNA control is used as negative control. 48 hours post transfection, cells were harvested for qRT-PCR for the analysis of VEGFA and BCL2 (N=3) (D) HUVECs were transfected with 30nM of circNPHP1 siRNA, miR-221-3p mimic respectively or in combination. Combination of control siRNA and control mimic is used as negative control. 48 hours post transfection, cells were harvested for qRT-PCR for the analysis of VEGFA and BCL2 (N=3). In (C-D) fold change in mRNA expression is relative to negative control; 18S is used as housekeeping gene. Results in (A–B) were assessed by unpaired Student *t* test between control siRNA and circNPHP1siRNA among the three groups respectively and in (C-D) by 1-way ANOVA (Dunnett’s post hoc test). P-values are indicated accordingly, P>0.05 is indicated as ns [nonsignificant].

## DISCUSSION

Identifying the molecular defects which guide EC dysfunction in the human ischemic and diabetic heart could vastly improve the understanding of the pathology and consequently support the design and adoption of better preventive and curative treatment. RNA therapeutics are gaining momentum in experimental and clinical cardiology^8^. Targeting the non-coding genome offer new perspectives^17^, and a miRNA-targeting drug has reached phase-II testing^9^ in cardiology. This anti-fibrotic drug represents the first successful attempt to reach to this advanced stage of translation of any noncoding RNAs (ncRNA). Despite initial promises^50,51^, vascular therapies based on miRNA repression have not yet successfully proceeded through the sequential layers of in-human trialling. CircRNAs constitute an endogenous control mechanism capable to specifically regulate the posttranslational activities of miRNAs. Targeting circRNAs to modulate miRNAs activities represent a testable alternative to the direct miRNA targeting by lock nucleic acid (LNA) or other oligonucleotide-based drugs. However, to the best of our knowledge, therapies targeting biological circRNAs remain untested in humans and represent a research field ripe for fundamental discoveries with the potential to ignite translational progresses. Notably, the enhanced stability of synthetic circRNAs compared to linear mRNA forms is currently of considerable interest for the pharmaceutical industry^52^, but to the best of our knowledge, not yet exploited to the therapeutical regulation of miRNAs.

The interactions between different RNA species are largely unexplored in clinical-relevant models. We present here, for the first time, regulatory networks involving circRNAs, miRNAs and mRNAs in the human heart affected by IHD leading to surgical revascularization. As part of this study, we have obtained an atlas of circRNAs, miRNAs and mRNAs present in the human left ventricle of patients suffering of severe IHD associated or not with T2DM and in a control surgical group represented by non-IHD, non-diabetic patients operated for repairing a mitral valve not compromised by a rheumatic aetiology. The data have been obtained and analysed as part of an observational prospective clinical study in cardiac surgery, expressively dedicated to noncoding RNA analyses in IHD and T2DM.

To guarantee the rigour of the work, the plan of analyses was designed and shared with the ethic committee ahead of accessing the RNA-sequencing data. Specifically, we identified circRNAs from whole transcriptome sequencing by putting together a pipeline using STAR and CIRCexplorer to detect back-spliced junctions. Several circRNA identification and annotation tools are currently available [reviewed in^18^]. For accurate identification of circRNAs from sequencing data, it is important to reduce false-positive rate of detection of back spliced junction reads. This can be achieved by considering detection of these junctions in several samples to gain more confidence or by applying a strict threshold on number of reads mapping to these junctions^53^. Most circRNA identification tools depend on aligners such as Bowtie^54^, BWA^55^ and STAR^27^. Several circRNA identification programs are available including Find_circ (https://github.com/marvin-jens/find_circ), CIRI^56^, CIRCexplorer^31^, CircTest ^57^, KNIFE^58^ etc. Among these, CIRCexplorer and KNIFE achieved the best sensitivity in detecting circRNAs^59^ which is why we incorporated it in our detection pipeline. We also identified mRNAs and miRNAs from our whole transcriptome and small RNA sequencing data respectively. Following detection of the various RNA species and performing differential expression (DE) analysis, we used streamlined bioinformatics approaches to construct circRNA-miRNA-mRNA networks in IHD and IHD+T2DM with a particular focus on EC biology and processes mediating microvascular disease and repair (apoptosis, proliferation, angiogenesis). Our study provides unique information on the putative interactions between these three RNA species (circRNA, miRNA and mRNA) in IHD with/out T2DM and is a useful resource for developing testable hypothesis for future therapeutic studies. Our study highlights a novel proangiogenic subnetwork, possibly activated to counteract the ischemic damage by growing new microvessels, which is commanded by circNPHP1.

The role of circRNAs as sponges of miRNAs and their implication in heart and circulatory disease has been studied, mostly using animal models (revised in^15,17^).

A recent study predicted the circRNA-miRNA-mRNA interactions in the human post-myocardial heart at an advanced stage of dilation and failure^24^. In this study, the LV samples were obtained from either the non-infarcted remote myocardium, during surgical ventricular reconstruction procedure, or from hearts explants derived from transplantation procedures^24^. The cell biology functional analyses were restricted to investigating circBPTF as regulator of cell death in HUVECs^24^. Previous studies have demonstrated the associations between T2DM and circRNAs in diabetic cardiomyopathy in animal models. These include circRNA_000203 (MYO9A), circRNA_010567 (CDR1), and circHIPK3 which are increased in diabetic mice and promote myocardial fibrosis^60, 61, 62^. We checked the levels of these circRNAs in our LV biopsy RNA-seq datasets and found circSLC8A1 and circHIPK3 to be increased in the LV of IHD with/out diabetes as compared to non-IHD patients whereas circMYO9A did not change and CDR1 was not detectable.

In the current study, circNPHP1 has been selected based on both expression in ECs and the predicted angiogenesis function of its commanded regulatory network encompassing downstream miRNA-mRNA partners. We next validated circNPHP1 to see whether it is upregulated and promotes the proliferation of both HUVECs and HCMECs kept under normal and disease mimicking conditions. CircNPHP1 did not influence apoptotic death. This suggests the prevalently reparative potential of circNPHP1 in the heart affected by CAD-induced angina, when the support of improved microcirculation could help boost the arrival of oxygens to the myocytes. Additional cardiac functions of this ubiquitously expressed circRNA are expected and deserve future investigations. To date, circNPHP1 has not been studied in the context of cardiovascular physiology and disease. CircNPHP1 is derived from exons 9 and 10 of NPHP1 gene. Although other circular isoforms emerging from NPHP1of the circular RNA were identifiable in our RNA-seq datasets, their expression was scarce or represented in very few samples. The linear NPHP1 transcript was uniformly expressed in the healthy and diseased heart as evidenced from our RNAseq on heart LV biopsies and confirmed by single nuclei RNA-seq analyses, which indicated its expression to be ubiquitous, even if the binding partners and function might vary across different cell types. NPHP1 encodes for the protein *Nephrocystin 1* that was initially discovered to be localized to the primary cilia and apical surface of kidney epithelial cells and interacts with molecules that take part in cell adhesion, signalling and maintenance^63^. NPHP1 has been associated to kidney disease (nephronophthisis) caused by an autosomal recessive whole gene deletion resulting in abnormal structure and function of primary cilia^64^. In nephronophthisis patients, cardiac defects like septal and aortic valve anomalies have also been reported^65^. The direct involvement of NPHP1 in heart disease has not been reported to date and information from GWAS Catalog (EMBL-EBI; https://www.ebi.ac.uk/gwas/) indicates no involvement in IHD or T2DM. Worth noting is the fact that although the cardiac levels of NPHP1 do not vary either between CAD with/out T2DM and non-CAD patients (as from our bulk RNA-seq data) or between the human healthy and diabetic heart (as from single nuclei RNA-seq analyses), one of the circular isoforms of this gene (circNPHP1, studied here) does. Based on our bulk RNAseq data, circNPHP1 levels are significantly upregulated in CAD associated with T2DM as compared to either CAD alone or controls, while they do not differ significantly between CAD and control groups. However, qRT-PCR analysed detected increased circNPHP1 in CAD

either associated or nor with diabetes vs control. This might reflect technical differences, particularly for data normalization, between the two technologies. The findings of LV-qRT-PCR are in line with data collected from ECs cultured under disease mimicking conditions. The linear form of NPHP1 remains unchanged between groups, as also indicated by single nuclei data in ECs resident in diabetic and non-diabetic heart donors. We also checked expression of circNPHP1 and linear NPHP1 in patient plasma samples from IHD with/without T2DM and non-IHD patients. The expression of circNPHP1 was not detectable across any the groups, whereas linear NPHP1 exhibited very low expression (Supplementary Figure 7). This indicates low release of both the forms from heart into plasma.

We found endogenous circNPHP1 to promote both EC proliferation and EC networking in capillary-like cords under hypoxia and HG (used to reflect the disease conditions in the dish). Other circRNAs were already reported to modulate endothelial function ^66,67,68^. However, in the context of human CAD leading to IHD, the presence, interactions and functions of endothelial circRNAs remain largely unexplored.

Using our streamlined bioinformatics pipeline involving integration of different types of data including expression profiles of circRNA, miRNAs, mRNAs, published EC data, predictions on possible interactions between the three, mapping of biological processes and pathways, we extracted a novel proangiogenic subnetwork commanded by circNPHP1 along with its predicted sponged miRNA partners (n=9) and their downstream mRNA targets (n=38). To confirm the interaction of the identified miRNAs with circNPHP1, pulldown of circNPHP1 was carried out in HUVECs by using antisense oligo probes (probe 1 and 2) designed against the sequence of its backsplice junction^44,69^. Among the predicted miRNAs, we found only miR-221-3p to be effectively enriched. Although we predicted miR-222-3p as well which is highly homologous to miR-221-3p and encoded in tandem from a gene cluster and often operating together^70^; we did not observe enrichment in miR-222-3p. Although the pulldown of circNPHP1 with probe 1 was effective in the ECs, the enrichment of miR- 221-3p with probe 1 was poor unlike significant enrichment with probe 2. This could be explained considering that probe 1 has a larger size which might bind other factors and can cause steric hindrance in binding of miR-221-3p. Additionally, circRNAs can possess certain conformational states which might confer differential binding to various factors^71^.

In the proangiogenic subnetwork of circNPHP1, miR-221-3p was found to interact with BCL2 and VEGFA, which are both involved in angiogenesis^72,73^ and BIM (BCL2L11) that works together with BCL2 in a context-dependent manner. miR-221-3p expression was already reported to be elevated in the rat post-MI^46,70^. The antiangiogenic role of miR-221-3p was previously demonstrated and explained with its repression of hypoxia inducible factor-1α (HIF-1α)^46^. Additionally, miR-221-3p targeting of Angiopoietin-2 was reported to impair the proliferation and cord formation capacities of hypoxic HCMECs and to inhibit the formation of tip cells, which play a vanguard role in angiogenesis^48^. In apparent contrast with these and our own data, miR-221-3p was proposed to promote proangiogenic and wound healing properties exerted by endothelial progenitor cell-derived exosome and to increase VEGF expression^74^. VEGF-A has long been established to play a therapeutic angiogenic role in experimental models of MI^75–77^. After several unsuccessful VEGF gene therapy clinical trials to induce regenerative angiogenesis in patients with cardiovascular disease^78^, a VEGF-A protein therapy by a naked modified mRNA without lipid encapsulation (AZD8601), co-developed by Astra Zeneca and Moderna successfully passed a randomized phase I study in otherwise healthy volunteers with T2DM (NCT02935712) (NCT02935712^79,80^) and next entered a randomized, double-blind, placebo-controlled multicentric safety study (EPICCURE) in IHD patients with moderately decreased left ventricular function (ejection fraction 30%–50%) undergoing elective CABG^78^ (this population is similar to the ARCADIA IHD groups). AstraZeneca and Moderna discontinued the investment after announcing that the Phase 2a study met the primary endpoint of safety and tolerability and that exploratory outcomes indicated potential improvement in cardiac composition and function^81^, even if the study was very limited in size (n=11 patients in total, for the 2 study groups study). Notwithstanding, these translational efforts have confirmed both the potential of promoting the local increase in soluble VEGF-A in the ischemic heart and the validity of non-viral delivery approaches. As discussed above, therapeutic circRNAs delivered via LNP could open new possibilities for improving the time-controlled regional increased of VEGF-A and other cardioprotective and regenerative proteins via microRNA sponging. Specifically, to circNPHP1, BCL2 could be elevated concomitantly with VEGF-A.

BCL2 also plays a crucial role in regulating the mitochondrial oxidative stress and angiogenesis in IHD^73^ and rat heart failure^82^. Our results in ECs confirmed that endogenous circNPHP1 dictates VEGFA and BCL2 expression by sponging miR-221- 3p. By contrast, BIM which is largely pro-apoptotic^83,84^ was unaffected upon modulation of circNPHP1.

We further investigated whether the circNPHP1/miR-221-3p axis is operational in other cell types. We also checked the binding of circNPHP1 to miR-221-3p in human AC16-cardiomyocyte and cardiac fibroblasts. The expression of linear and circNPHP1 in AC16 is comparable to that of the ECs, while cultured cardiac fibroblasts showed low expression of circNPHP1. Regardless, circNPHP1 KD reduced the proliferation of cardiac fibroblasts. By contrast, circNPHP1 KD did not affect AC16 proliferation or apoptosis, suggesting that it might regulate other phenotypes. miR-221-3p could be co-precipitated with circNPHP1 in both AC16 and fibroblasts, suggesting that the circNPHP1/miR-221-3p axis is potentially operational in different cardiac cell types and the need for further investigation.

## STUDY LIMITATIONS

The mechanistic part of this study has focussed on EC and angiogenesis. Notwithstanding, the ARCADIA RNA-seqs allow for expanding this focus in multiple directions. Our datasets are available for further analyses by us and others and are expected to expand beyond the specific focus kept in this first study. *In vivo* assessment of the pro-angiogenic subnetwork with regards to cardiac functions and reparative angiogenesis in animal models remain to be explored.

## CONCLUSION

In conclusion, our study has identified novel differential circRNA-miRNA-mRNA networks in IHD and IHD+T2DM with a particular focus on ECs. It specifically highlights the proangiogenic axis involving circNPHP1/miR-221-3p/VEGFA/BCL2 in

IHD and IHD+T2DM implicating its potential significance for therapeutic angiogenesis and cardiac repair.

## CLINICAL PERSPECTIVES

### COMPETENCY IN MEDICAL KNOWLEDGE

CAD-induced IHD is a major cause of mortality and often worsened by T2DM. Coronary EC function is highly compromised in IHD associated with T2DM. Elucidation of novel mechanisms of gene expression regulating the cardiac endothelium is of high significance for therapy. CircRNAs are critical regulators of gene expression thereby controlling various cellular processes. This is the first time that circRNAs and their networks have been mapped in IHD and T2DM using clinical samples from left ventricle biopsies of cardiac surgery patients. We have elucidated a novel pro-angiogenic arc (circNPHP1/miR-221-3p/BCL2/VEGFA) that regulates angiogenesis and proliferation in cardiac endothelial cells.

### TRANSLATIONAL OUTLOOK

Our study highlights a potential protective role of circNPHP1 in reparative angiogenesis in IHD and T2DM and can be explored further as a novel therapeutic target.

Delivery of circNPHP1 via LNP could be of significant therapeutic potential for regional increased of proangiogenic pathways, offering a new strategy for vascular regeneration in ischaemic heart disease.

## DATA AVAILABILITY

Following acceptance of the paper for publication, the ARCADIA human left ventricle RNA-seq datasets employed in this study will be published at NCBI-GEO database (https://www.ncbi.nlm.nih.gov/geo/). Additionally, the data created and/or analysed in the current study will be made available through online repositories.

## ETHICS APPROVAL AND CONSENT TO PARTICIPATE

All human samples were obtained in accordance with the principles of the Declaration of Helsinki. The study was reviewed and approved by the National Research Ethics Service Committee London–Fulham (date of approval, 20 Dec 2013; reference REC 13/LO/1687).

## AUTHOR CONTRIBUTIONS

MA and MS contributed equally. MA designed and conducted the bioinformatics part of the study, in line with the predefined plan for analyses of the ARCADIA study. She stored and classified the data (together with MS) and wrote an advanced draft of the manuscript (together with MS), preparing all the bioinformatic illustrations. MS performed the vast majority of cell and molecular biology experiments, stored and classified the data (together with MA) and wrote an advanced draft of the manuscript (together with MA), preparing the figures emerging from laboratory work. KF contributed to store the clinical samples and to write ethical documents. She optimised and performed the RNA extraction and quality control of the ARCADIA samples; GDA obtained research funds and contributed to design and conduct the ARCADIA study; PPP obtained research funds and contributed to conduct the ARCADIA study, AL collected and processed clinical samples and conducted parts of the laboratory experiments; ACJ contributed to design and supervise parts of the laboratory experiments; JJ and PKS performed the analyses on the single cell RNA-seq datasets. EP contributed to design the plan for analyses of the ARCADIA study and supervise part of the bioinformatic analyses; CE obtained research funds, contributed to design the ARCADIA study, and wrote it. She designed the cell biology work together with MS and wrote the final version of the manuscript with MA and MS. All authors read and approved the submitted manuscript.

## ACKNOWLEDGEMENTS

We are grateful to Dr. Fabio Martelli and Dr. Alessia Bibi for critical discussions and advice regarding the circRNA pulldown experiments. We are also thankful to Prof. Pilar Ruiz-Lozano, Dr. Hassan Abdulrazzak and Junqing Zhang for providing HCMECs and human cardiac fibroblasts; Dr. Sharadkumar Kholia, Dr. Walid Sweaad, Dr. Pragati Panday, Junyi Liu and Casey Lau for general support with this study. Dr Cristina Beltrami supported us in the initial phase of the ARCADIA sample collection. Panagiotis Kyriazis (Imperial College) and several Bristol clinical trial unit staff supported us for the administrative aspects of the ARCADIA study. We are also grateful to the patients, surgeons and research nurses and Bristol clinical trial unit staff who contribute into the ARCADIA studies. For illustrations, we used some images from Vecteezy (www.vecteezy.com).

## SUPPLEMENTARY DATA

**Supplementary File 1: ARCADIA Study Plan**

**Supplementary File 2: Resources table**

**Supplementary Table 1: Patient characteristics ARCADIA study**

