## Supplementary figures for "Human left ventricle circRNA-miRNA-mRNA network analyses reveals a novel proangiogenic role for circNPHP1 under ischemic conditions"

#### Supplementary Figure 1

A. CircRNA identification workflow

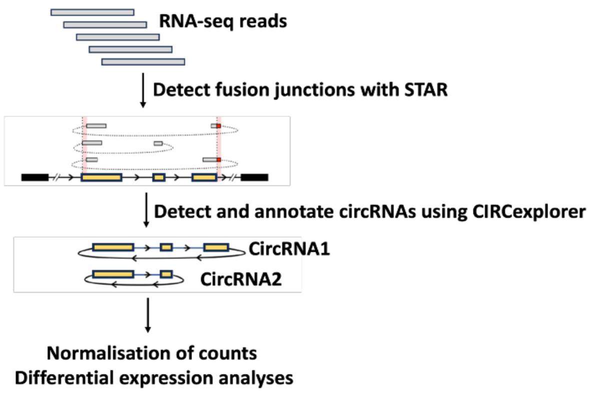


B. Numbers of detected circRNAs

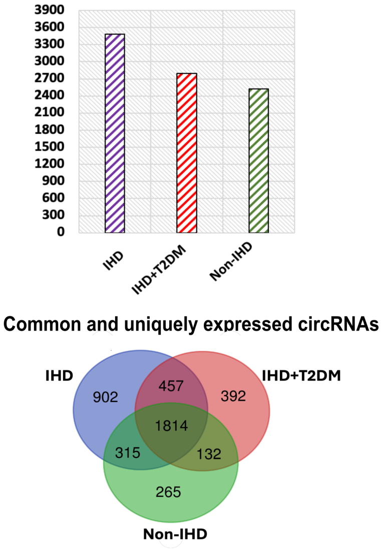


C. No. of significant differentially expressed circRNAs, miRNAs and mRNAs

| Comparison | circRNAs | miRNAs | mRNAs |
|---|---|---|---|
| IHD vs Non-IHD | 850 | 35 | 2824 |
| IHD+T2DM vs Non-IHD | 585 | 18 | 2266 |
| IHD+T2DM vs IHD | 511 | 48 | 431 |

D.

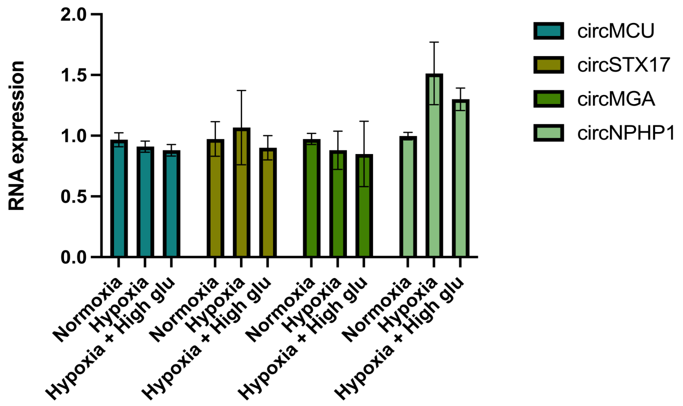


#### Supplementary Figure 1: Identification and quantification of circRNAs

(A) *CircRNA identification workflow:* RNAseq reads from whole transcriptome were first aligned using STAR aligner then circRNAs were identified using CIRCexplorer with annotations from CircBase. Following that, raw counts were normalised to

Fragments per Million (FPM) and differential analyses was carried out using R package *Limma.*
(B) Bar graph showing number of detected circRNAs in IHD, IHD+T2DM and non-IHD. CircRNAs that had non-zero expression values in at least 50% of the samples of a patient group were considered expressed/detected in that particular group. Venn diagram shows common and uniquely expressed/detected circRNAs in the three patient groups: [IHD(N=12), IHD+T2DM(N=11), non-IHD(N=12)].
(C) Number of significant differentially expressed circRNAs, miRNAs and mRNAs in LV biopsies (log2 FC threshold of +/-0.58 and P-value < 0.05); [IHD(N=12), IHD+T2DM(N=11), non-IHD(N=12)].
(D) Preliminary screening of circRNAs: HCMECs were cultured in normal conditions, hypoxia (1% Oxygen) and hypoxia-high glucose (High Glu, 25mM D-glucose) conditions respectively. After 48 hours, cells were harvested for qRT-PCR for the analysis and validation of indicated circRNAs. RNA expression is relative to normoxia; 18S is used as housekeeping gene (N=1, technical replicates: n=3).

#### Supplementary Figure 2

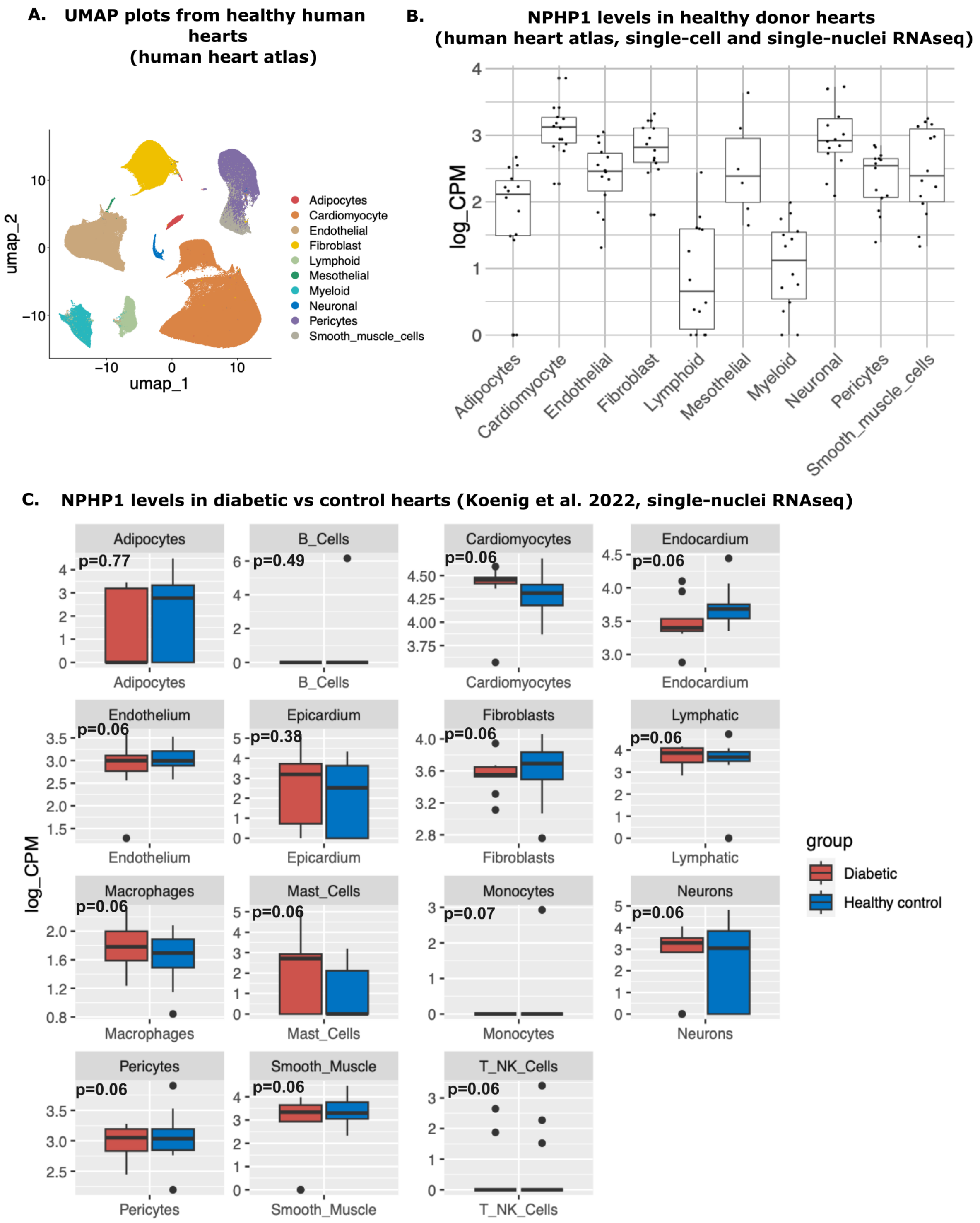


#### Supplementary Figure 2: Linear NPHP1 expression in heart cells

(A) Single-cell and single-nuclei RNAseq data from healthy donor hearts (human heart atlas) was used to make UMAP plots showing data in different cell types. (B)

Expression levels of NPHP1 gene as log2 transformed counts per million (CPM). Each dot indicates the data point as one donor (source: human heart atlas). (C) Single-nuclei RNAseq data of control hearts from Koenig et al (2022). Among 26 donors used as controls in this study, 9 donors had diabetes. Boxplot showing expression levels of NPHP1 gene as log2 transformed counts per million (CPM) in diabetic and healthy controls, each dot indicates the data point as one donor. Wilcoxon test was performed to calculate P-values. Differences between diabetic and healthy patients were considered significant if they passed the p-value threshold of less than 0.05.

#### Supplementary Figure 3

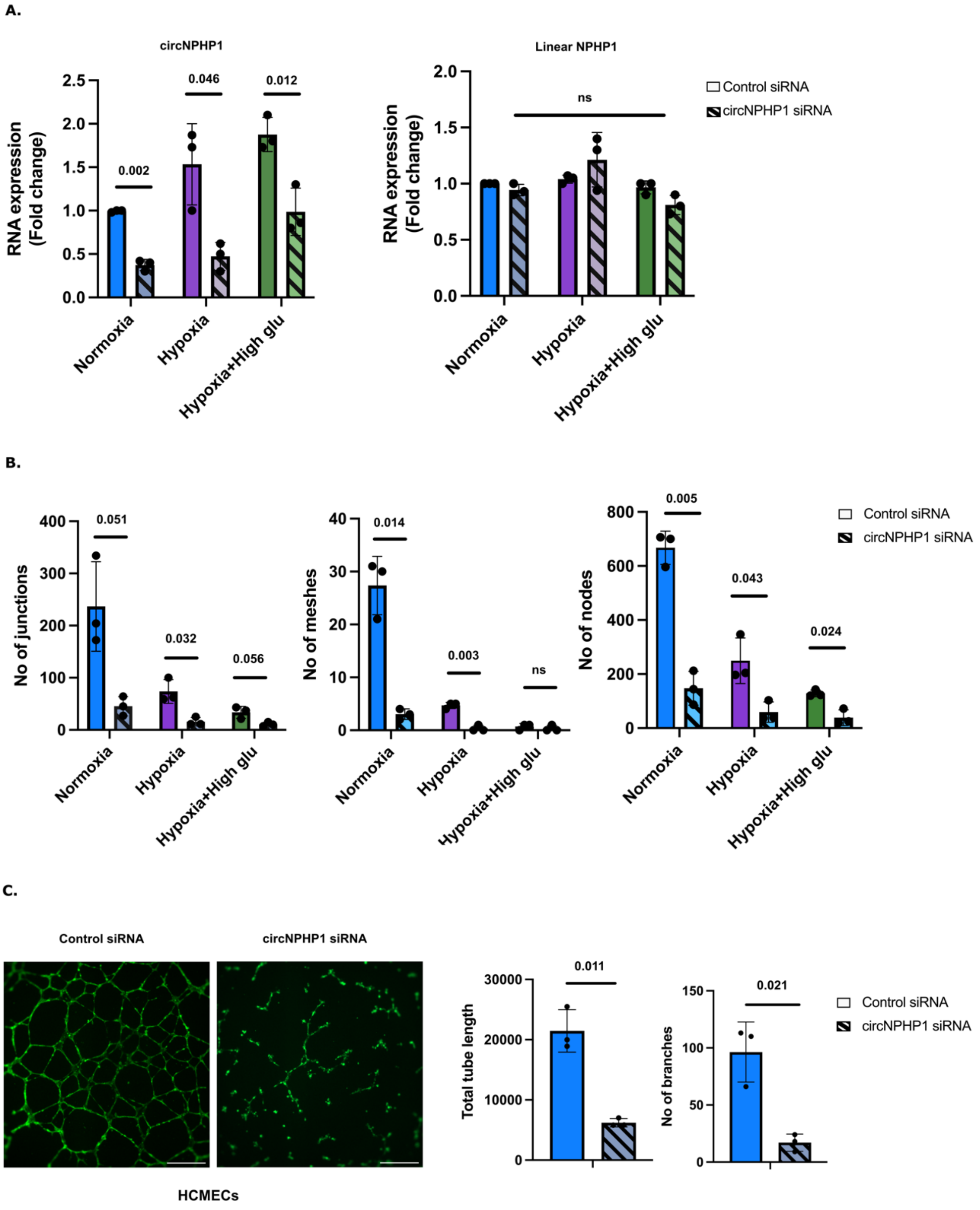


**Supplementary Figure 3: CircNPHP1 is involved in proliferation and angiogenesis in the ECs**

(A) HCMECs were transfected with 30nM of circNPHP1 short interfering RNA (siRNA) or control siRNA, and cultured in normal conditions, hypoxia (1% Oxygen) and hypoxia-high glucose (25mM D-glucose) conditions respectively. 48 hours post transfection, cells were harvested for qRT-PCR for the analysis of circNPHP1 (left panel) and linear NPHP1 (right panel) (N=3). Fold change in RNA expression is relative to control siRNA; 18S is used as housekeeping gene.

(B) HUVECs were transfected with 30nM of circNPHP1 siRNA or control siRNA, and cultured in normal conditions, hypoxia (1% Oxygen) and hypoxia-high glucose (25mM D-glucose) conditions respectively. 48 hours post transfection, cells were seeded on a 96-well plate containing Growth Factor Reduced Matrigel and cultured in similar conditions for 8 hours to determine cord formation. Histograms depicting the total number of meshes, junctions and nodes quantified from the angiogenesis assay (Figure 5B) (N=3).

(C) HCMECs were transfected with 30nM of circNPHP1 siRNA or control siRNA. 48 hours post transfection, cells were seeded on a 96-well plate containing Growth Factor Reduced Matrigel for 8 hours to determine cord formation. Representative images of angiogenesis (stained with Phalloidin, scale bar: 500 mM) (left panel). Histograms depicting the total tube length and number of branches quantified from the angiogenesis assay (right panel) (N=3).

Results in (A–C**)** were assessed by unpaired Student *t* test between control siRNA and circNPHP1siRNA among the three groups respectively. P values are indicated accordingly, P>0.05 is indicated as ns [nonsignificant].

### Supplementary Figure 4

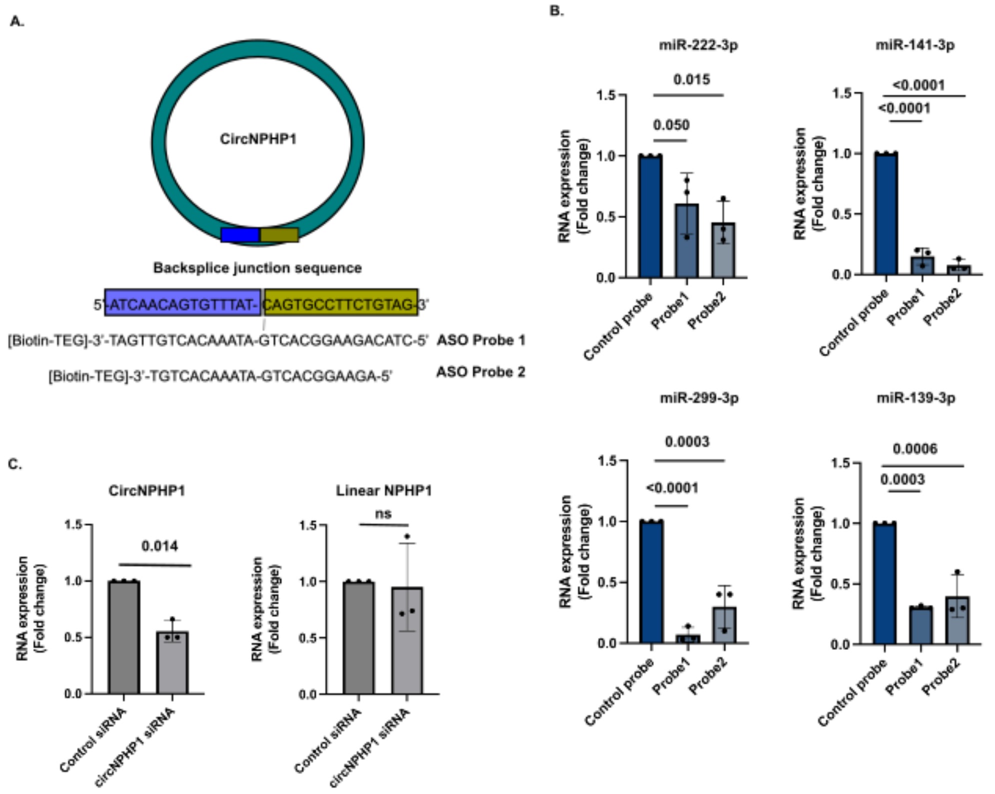


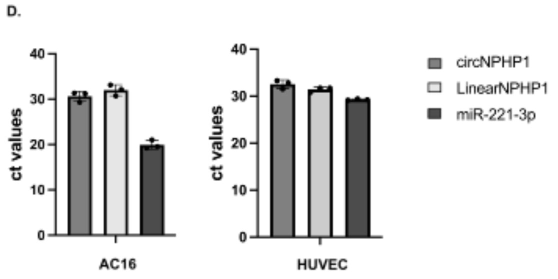


| Human cardiac fibroblasts | Mean Ct values |
|---|---|
| circNPHP1 | 34.4 |
| Linear NPHP1 | >35 |
| 18S | 12 |
| miR-221-3p | 25.2 |
| U6 | 25.9 |

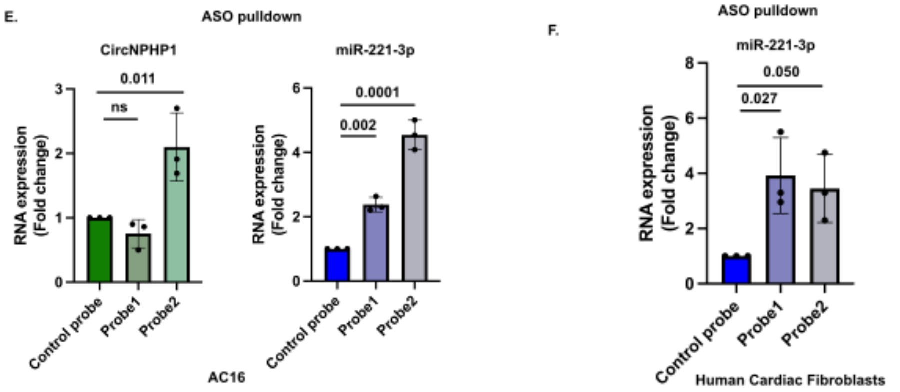


| Mean Ct values | miR-221-3p | U6 | circNPHP1 | 18S |
|---|---|---|---|---|
| Control probe | 34.8 | 32.5 | >35 | 16 |
| Probe 1 | 31 | 31.3 | >35 | 12.1 |
| Probe 2 | 31.1 | 30.5 | >35 | 11.9 |

Human Cardiac Fibroblasts

**Supplementary Figure 4: CircNPHP1 binds miR-221-3p and constitutes a pro-angiogenic sub-network**

(A) Schematic representation and sequence of the biotinylated probes used for pulldown of circNPHP1. Antisense oligo (ASO) probe 1 was designed by joining the last 15 nucleotides of circNPHP1 (green) to the first 15 nucleotides (blue) to make a 30-nucleotide sequence against the backsplice junction. The second probe (probe 2) consists of 22 nucleotide sequence against the backsplice junction, which is also the siRNA sequence for circNPHP1 used in this study.

(B) HUVECs extracts were incubated with biotinylated control probe, probe 1 and probe 2 respectively (details in methods) for pulldown of circNPHP1 using streptavidin beads, followed by RNA isolation and qRT-PCR analysis of miR-222-3p, miR-141-3p, miR-139-5p, miR-299-3p. Fold enrichment was calculated in the pulldown samples with probe 1 and probe 2 against the control probe pulldown. 18S and U6 were used as reference genes to normalize for circNPHP1, linear and the miRNAs respectively (N=3) (Refer Figure 6B).

(C) HUVECs were transfected with 30nM of circNPHP1 short interfering RNA (siRNA) or control siRNA. 48 hours post transfection, cells were harvested for circNPHP1 pulldown (**Refer figure 6C**) qRT-PCR for the analysis of circNPHP1 (left panel) and linear NPHP1 (right panel) (N=3) in the input samples. Fold change in RNA expression is relative to control siRNA; 18S is used as housekeeping gene.

(D) Histograms depicting the cycle threshold (ct) values from qRT-PCR of circNPHP1, linear NPHP1 and miR-221-3p (N=3) in AC16, HUVEC and cardiac fibroblasts (table). Ct values > 35 were considered undetectable.

(E) AC16 (F) cardiac fibroblast extracts were incubated with biotinylated ASO control probe, probe 1 and probe 2 respectively (see details in methods) for pulldown of circNPHP1 using streptavidin beads. Subsequently, RNA was isolated and subjected to qRT-PCR for the analysis of circNPHP1 (left panel) and miR-221-3p (right panel) (N=3). Fold enrichment was calculated in the pulldown samples with probe 1 and probe 2 against the control probe pulldown. 18S and U6 were used as reference genes to normalize for circNPHP1 and miR-221-3p respectively (N=3).

The table in (F) shows the average Ct values from pulldown in cardiac fibroblasts. Ct values > 35 were considered undetectable. Biological replicates in AC16 and cardiac fibroblast are carried out on cells from different aliquots and passages of the same lot.

Results in (B, E and F) were assessed by 1-way ANOVA (Dunnett's post hoc test) and in (C) by unpaired Student *t* test. P values are indicated accordingly, $P>0.05$ is indicated as ns [nonsignificant].

#### Supplementary Figure 5

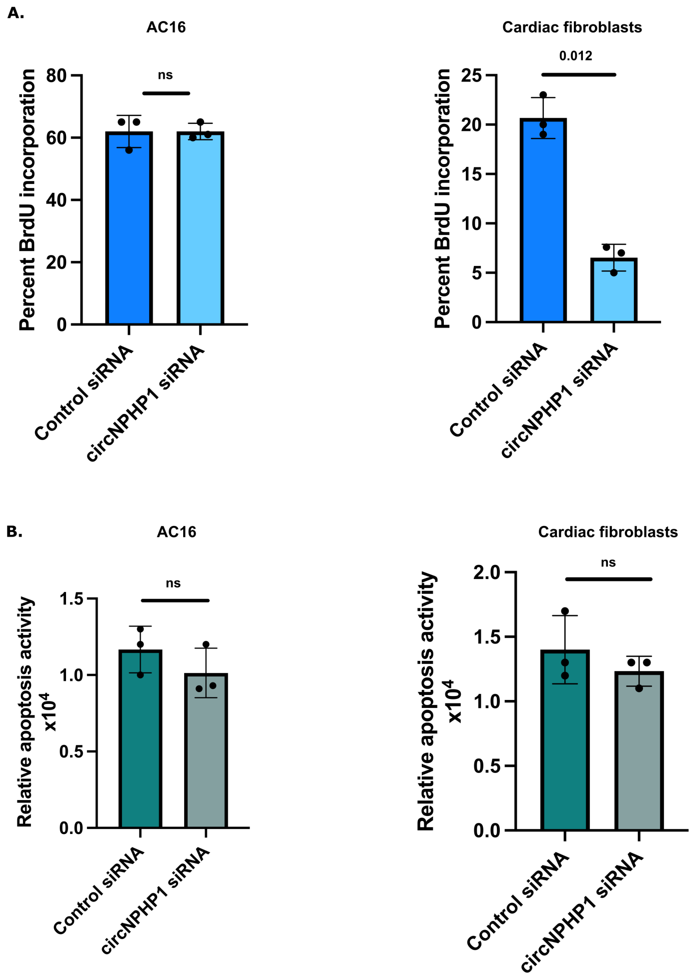


**Supplementary Figure 5: CircNPHP1 regulates cellular proliferation in cardiac fibroblasts**

(A) AC16 (left panel) and cardiac fibroblasts (right panel) were transfected with 30nM of circNPHP1 siRNA or control siRNA. 48 hours post transfection, cells were harvested for proliferation assay (BrDU incorporation) **(N=**3) and (B) apoptosis assay (**N=3)** . Results were assessed by unpaired Student *t* test. P values are indicated accordingly, P>0.05 is indicated as ns [nonsignificant]. Biological replicates in AC16 and cardiac fibroblast are carried out on cells from different aliquots and passages of the same lot.

**Supplementary Figure 6**

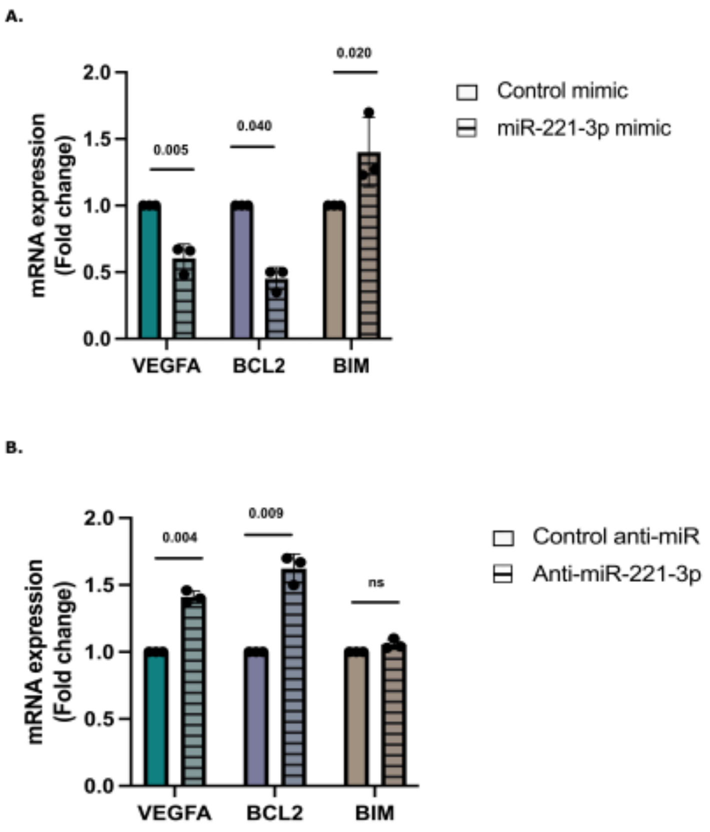


**Supplementary Figure 6: miR-221-3p regulates VEGFA and BCL2**

(A) HUVECs were transfected with 30nM of miR-221-3p mimic or control mimic (B) anti-miR-221-3p or anti-miRNA negative control. 48 hours post transfection, cells were harvested for qRT-PCR of BIM, BCL2 and VEGFA. Fold change in mRNA expression is relative to control (control mimic (A) or anti-miRNA negative control (B); 18S is used as housekeeping gene; N=3. Results were assessed by unpaired Student *t* test between the two groups among the three genes respectively. P values are indicated accordingly, P>0.05 is indicated as ns [nonsignificant].

**Supplementary Figure 7**

| | | Mean Ct values | | |
|---|---|---|---|---|
| | circNPHP1 | Linear NPHP1 | GAPDH | cel-miR-39 |
| **Non-IHD** | >35 | 33.06 | 32.7 | 30.71 |
| **IHD** | >35 | 33.56 | 32.25 | 30.67 |
| **IHD+T2DM** | >35 | 33.98 | 32.27 | 30.85 |

**Supplementary Figure 7: Quantification of the expression levels of CircNPHP1 and linear NPHP1 in the plasma of patients**

qRT-PCR analysis of circNPHP1 and linear NPHP1 of patient plasma from non-IHD (N=5), IHD (N=6), IHD+T2DM (N=4). GAPDH is used as endogenous control and cel-miR-39 as spike-in control.

The table shows the average Ct values. Ct values > 35 were considered undetectable.

**Supplementary Figure 8**

Uncut western blot: (Refer Figure 7B)

A. (representative image used in figure 7B)

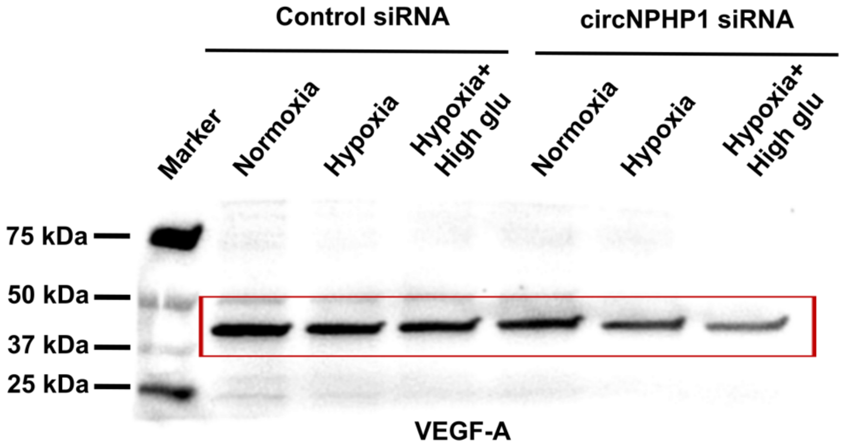


VEGF-A

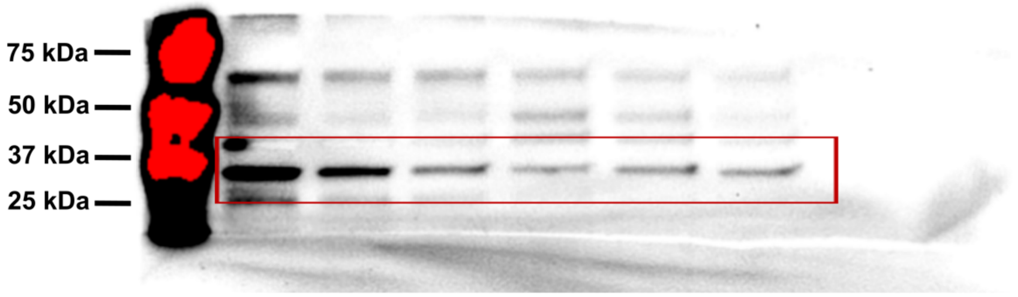


BCL2

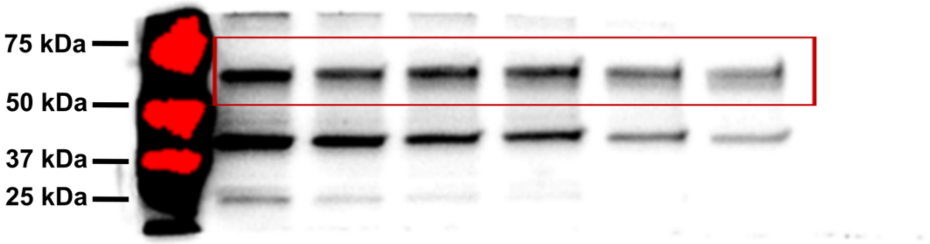


Lamin B1

B.

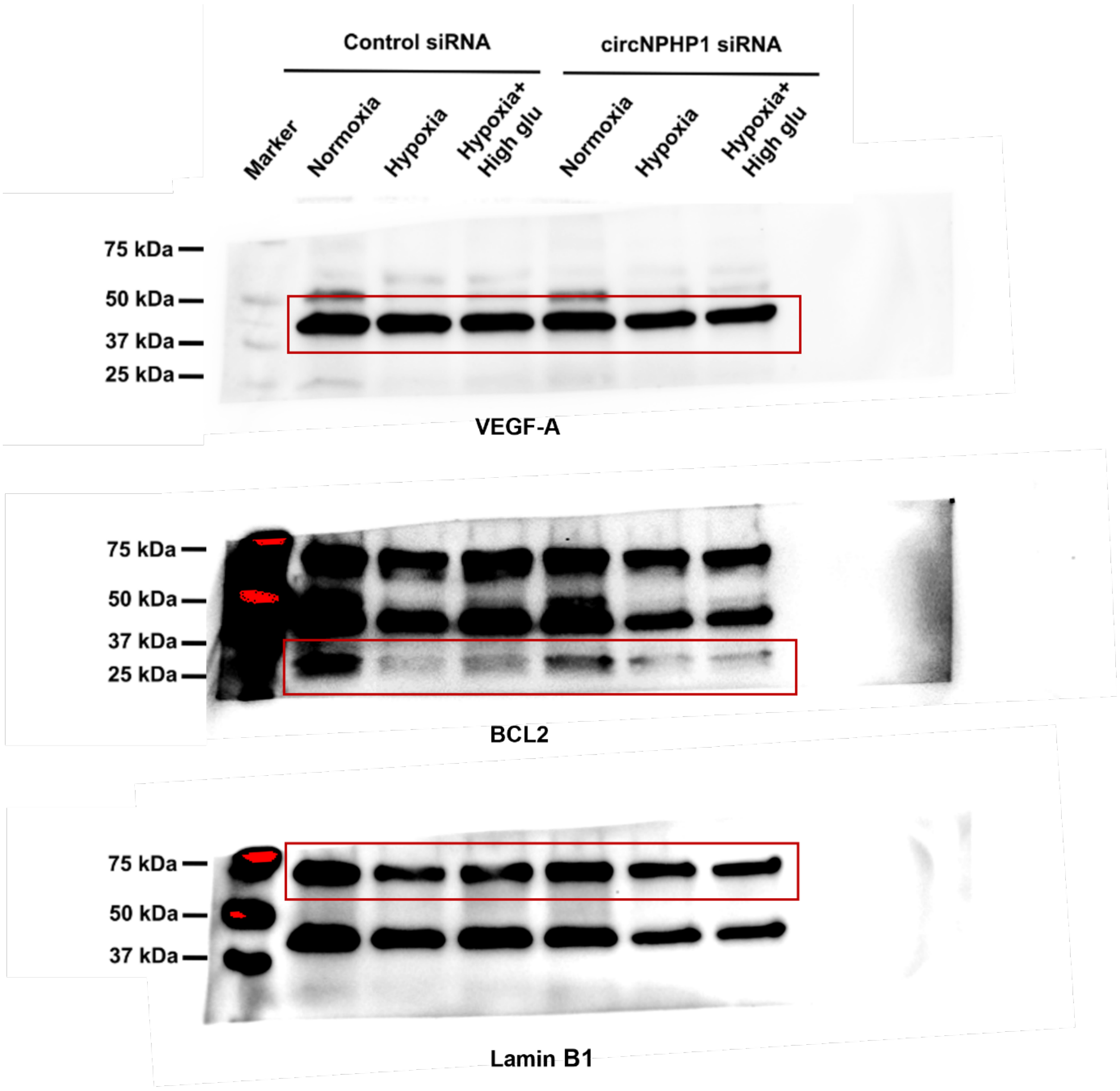

Control siRNA
circNPHP1 siRNA
Marker
Normoxia
Hypoxia
Hypoxia+ High glu
Normoxia
Hypoxia
Hypoxia+ High glu
75 kDa
50 kDa
37 kDa
25 kDa
VEGF-A
75 kDa
50 kDa
37 kDa
25 kDa
BCL2
75 kDa
50 kDa
37 kDa
Lamin B1

C.

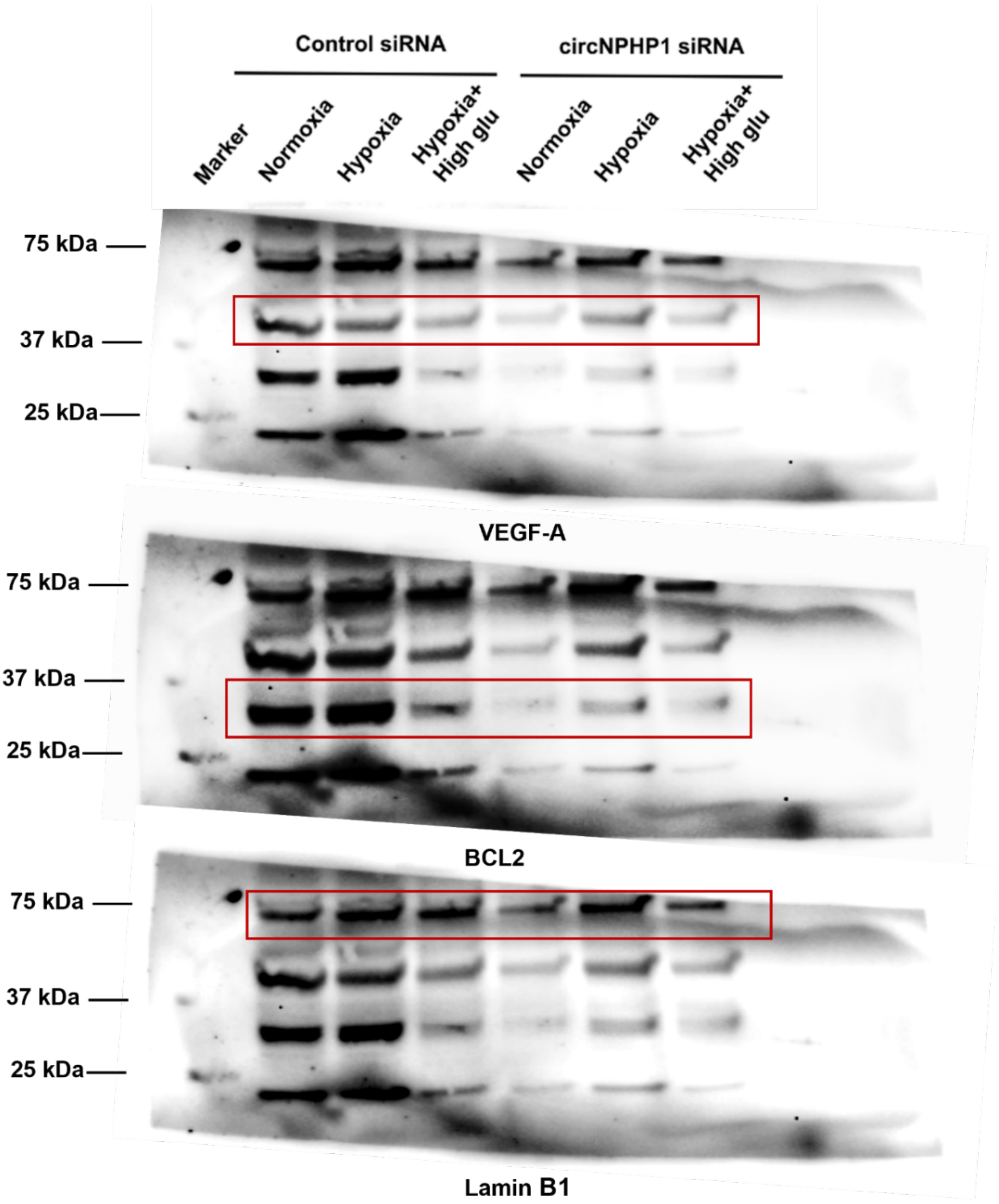


**Supplementary Figure 8** Uncut western blots with additional replicates used for **Figure 7B**

(A) Uncut blot of representative image used in figure 7B (B) and (C) are additional biological replicates.
