## Supplementary file 1 for "Human left ventricle circRNA-miRNA-mRNA network analyses reveals a novel proangiogenic role for circNPHP1 under ischemic conditions"

Bristol Medical School
University of Bristol
Research Floor Level 7
Bristol Royal Infirmary
Upper Maudlin Street
Bristol, BS2 8HW

London Fulham Research Ethics Committee
Charing Cross Hospital
London, W6 8RF

4th January 2018

To whom it may concern,

Re: ARCADIA stage 1 analysis plan v1.0

We are sending this document in advance of doing the proposed analyses described therein, to show that these analyses were pre-planned, thus minimising the risk of bias from subsequent selection of analyses or results.

For this document to fulfil its function, it is imperative that it is acknowledged and date-stamped by an independent organisation, hence we are sending it to the REC. Please acknowledge receipt of this document and file it with the other documents relating to this study (REC reference 13/LO/1687).

Yours faithfully,

Professors Gianni Angelini (Chief Investigator), Costanza Emanueli (Co-Investigator and Bristol Principal Investigator), Enrico Petretto (Principal Investigator), Barnaby Reeves (PI, Epidemiologist and Trialist)

**ARCADIA planned analyses – stage 1 samples**

**REC reference: 13/LO/1687**

This document has been sent to the REC in advance of doing the proposed analyses, to show that the analyses described in the document were pre-planned, thus minimising the risk of bias from subsequent selection of analyses or results.

### 1. Background

In this research programme, we aim to identify ncRNAs and related pathways affected by ischaemic heart disease (IHD) and type 2 diabetes mellitus (T2DM), and integrate this with data from proteomics and metabolomics studies. We have 4 patient study groups:

1. Patients with IHD ungergoing CABG surgery (CABG-NDM)
2. Patients with IHD undergoing CABG surgery, with T2DM (CABG-DM)
3. Patients undergoing AVR surgery
4. Patients undergoing MVR surgery

By comparing these patient groups we will consider the effects of 3 disease features: ischaemic heart disease (IHD), type 2 diabetes mellitus (T2DM), hypertrophic remodelling (RM). The following table indicates which features are present (1) or absent (0) in each group:

| Group | HD | T2DM | RM |
|---|---|---|---|
| 1. CABG-NDM | 1 | 0 | 0 |
| 2. CABG-DM | 1 | 1 | 0 |
| 3. AVR | 0 | 0 | 1 |
| 4. MVR | 0 | 0 | 0 |

From each patient group we have left ventricular biopsies, pericardial fluid, and plasma taken from various locations (peripheral and transcoronary) and at various time points (pre-, during-, 24h and 48h post-surgery).

We will employ data modelling using systems biology approaches to provide a rich dataset of functionally coherent transcriptional signatures associated with ischaemic disease (groups 1 and 2 vs. groups 3 and 4), the effect of T2DM (group 1 vs group 2) and the effect of hypertrophic remodelling (group 3 vs group 4) in the human heart, which will pinpoint single ncRNAs/mRNAs as well as ncRNA-mRNA networks and identify potential network regulators termed "regulatory ncRNA hubs". We will also investigate networks of ncRNAs which are conserved between the myocardium and the blood and the impact of IHD and T2DM on them. Finally, we will integrate with proteomic data from the tissue, and proteomics and metabolomics data from the biofluids.

### 2. Summary of planned statistical analyses

#### A. Left ventricular biopsies

Unsupervised systems-biology analyses of LV biopsy RNA-seq data will be developed through two parallel analytical tasks followed by systematic *in silico* annotation of the results:

1) *Task 1* (single ncRNA/mRNA level analyses). For each expressed ncRNA and mRNA, we will carry out genome-wide differential expression (DE) analysis between: (a) CABG (study groups 1 and 2) and the AVR and MVR controls (study groups 3 and 4); (b) between non-diabetes and T2DM in CABG (study group 1 vs. 2) and (c) between hypertrophic and non-hypertrophic conditions (study group 3 vs. 4). For the single ncRNA/mRNA DE analysis we will use the Limma R package, which provides stable results even with a small number of replicates

by borrowing information across all ncRNAs and mRNAs. False Discovery Rate (FDR) correction (q-value) will be employed to identify robust differentially expressed ncRNAs and mRNAs.

2) *Task 2* (network level analyses). The DE analysis will be extended at the level of multiple coexpressed ncRNAs and mRNAs to identify differential ncRNAs/mRNAs networks across the conditions considered i.e. (a) CABG (study groups 1 and 2) and the AVR and MVR controls (study groups 3 and 4); (b) between non-diabetes and T2DM in CABG (study group 1 vs. 2) and (c) between hypertrophic and non-hypertrophic conditions (study group 3 vs. 4). We will take advantage of a new algorithm (Cross-Conditions-Cluster-Detection or C3D) for efficient and parameter-free detection of differential co-expression networks across multiple patients cohorts (*i.e.* networks of ncRNAs and mRNAs that are differentially associated with CAD or T2DM), assessing empirical P-values and FDR associated with the detection of differential networks. The power of the C3D algorithm to recover differential networks in small samples has been previously demonstrated (and published, Xiao *et al*. PloS Genetics 2014) by Petretto using extensive simulation studies and showed in in C3D-applications to real genome-wide RNA-seq data using as few as 6 samples per group.

3) *In silico* annotation. To prioritize functionally relevant pathways and thereby identify candidate ncRNA and mRNAs for further investigations in LV biopsies and in human cell models, we will systematically annotate the identified co-expression networks of ncRNAs and mRNAs using three independent approaches:
(i) functional enrichment analysis using Gene Ontology and KEGG pathways as well as the “Cardiovascular Gene Ontology Annotation Initiative” for high-quality annotation of ~2500 cardiovascular-relevant genes (Cardiovascular Gene Ontology gene list, http://www.ucl.ac.uk/functional-gene-annotation/cardiovascular);
(ii) cell-type specificity using Cten tool which exploits large data bases of gene expression signatures across many cell-types, and so provides information of cell-type specific expression patterns;
(iii) analysis of miR target genes in the set of coexpressed genes (i.e. networks identified in *Task 2*) using miRWalk (a comprehensive human and mouse database that provides information on predicted binding to putative target genes and take advantage of multiple miR-targets prediction algorithms. To minimize false positives we will use only the ncRNA targets predicted by multiple algorithms. Then, for each network of co-expressed ncRNA/mRNAs we will employ out an *ad hoc* Gene Set Enrichment Analysis (GSEA) to: (i) prioritize networks enriched for co-expressed genes which are targeted by one or more miRs in human LV and (ii) pinpoint ncRNAs that regulate and are also co-expressed with mRNAs, therefore identifying potential “regulatory ncRNA hubs” of the “differential” networks associated with IHD and T2DM.

To take full advantage of the sequencing data generated, another outcome will be the identification of new RNA isoforms and modifications differentially expressed in the ischaemic human heart and regulated in T2DM. These will be identified by combining the EdgeR method with recently proposed hierarchical Bayesian multi-sample approaches, which have the advantage of borrowing information across all samples in estimating expression levels and improve the estimates for low abundance isoforms. The list of putative new RNAs and RNA modifications will be further analysed in collaboration with Dr. Ruth Lovering (UCL), who leads the aforementioned “Cardiovascular Gene Ontology Annotation Initiative” for highquality GO annotation of miR in cardiovascular systems.

**B. Pre- and peri-operative blood whole plasma**

This work aims to define the conservation of heart-derived RNAs in the peripheral circulation, therefore we wish to identify (i) RNAs and RNA networks which are produced and released by the myocardium in the coronary sinus blood and (ii) the myocardium-derived RNAs and RNA networks that are conserved in the peripheral plasma and which might therefore represent mediators of heart signals sent from the heart to different organs.

The plasma qPCR data will be compared to the RNA-seq data on LV biospies. This allows for calculation of the “transcoronary” RNA plasma concentration gradients, identifying RNAs which are either consumed or enriched through the myocardium passage and hence revealing RNAs that might be released (hence, likely produced) by the myocardial cells. To reinforce the evidence of their myocardial origin, the transcript forms of the RNA enriched in the coronary sinus will be also measured (by qPCR) in the LV biopsy. We will be able to establish the pool of RNAs expressed in the myocardium and released in the blood and the effect of IHD and T2DM on this. This will employ the parallel analytical approaches already described in **3A**.

First, we will carry out DE analysis at the single RNA level as well as the network level to identify differential RNAs and RNAs-networks associated with CAD and T2DM in plasma. Second, we will prioritize the "transcoronary" RNA patterns that are specifically conserved with the patterns identified in the LV biopsy. To confirm the myocardial origin of the "transcoronary" RNAs, these will be cross-matched with the RNAs previously identified in LV biopsies. To determine statistical significance, a permutation procedure will be adopted to assess the likelihood of observing conserved "transcoronary" RNAs signatures by chance and calculate their empirical P-value and FDR (this procedure is already implemented in the C3D algorithm). Only conserved "transcoronary" signatures with FDR<5% will be taken to the next step.

**C. Peri-operative blood plasma extracellular vesicles (EVs) – exosomes and microparticles (MPs)**
The same approach as in part **3B** will be taken on the EV fractions of the plasma.
We will use this to investigate the trafficking of the RNA *via* vesicles. RNA enrichment in these vesicles *vs*. the non-vesicular plasma component will be investigated and correlated to the RNA content in the LV biopsies.

**D. Pericardial fluid**
The same approach as in part **3B** will be taken on the pericardial fluid.

**E. Pericardial fluid extracellular vesicles (EVs) – exosomes and microparticles (MPs)**
The same approach as in part **3B** and **3C** will be taken on the pericardial fluid EVs.

3. <u>Summary of samples and laboratory methods of analysis</u>

In each of the sample sets and forms of laboratory analysis described in this section, comparisons will be made between groups as outlined in sections 1 and 2. The details of the statistical methods to be used in these comparisons can be found in section 2. In some cases, the planned sample sizes stated are not available due to missing samples. In such cases, we will adopt more stringent p-values to protect from inflated false positives.

**A. Left ventricular biopsies**

i. Whole transcriptome RNA sequencing (Imperial College London transcriptomics facility ): CABG-NDM, CABG-DM, MVR, AVR all samples
*(n = 12 per group; 48 samples total)*
ii. Small RNA sequencing (Exiqon/Qiagen - company based in Germany): CABG-NDM, CABG-DM, MVR, AVR all samples
*(n = 12 per group; 48 samples total)*
iii. qPCR validation of selected findings from sequencing: CABG-NDM, CABG-DM, MVR, AVR
*(n = 12 per group; 48 samples total)*
iv. Proteomics analysis (University of Bristol proteomics facility): CABG-NDM, CABG-DM, MVR, AVR all samples
*(n = 12 per group; 48 samples total)*

**B. Pre- and peri-operative blood whole plasma**

i. miRNA qPCR arrays (purchased from Exiqon): CABG-NDM, CABG-DM, MVR, AVR, pre-operative peripheral blood plasma and intra-operative transcoronary blood plasma
*(n = 3-5 samples per group*. One group has 3 samples, one group has 4 samples, and 2 groups have 5 samples each. For each group there are 3 plasma sample types: pre-operative, ascending aorta, coronary sinus.)*
ii. qPCR validation of ncRNAs involved in trafficking from LV (2Ai, Aii) to blood (2Bi): CABG-NDM, CABG-DM, MVR, AVR

*(n = planned 12 samples per group; 48 samples total, however it seems likely that some samples will be missing)*

iii. Time-course of expression in plasma of the ncRNAs identified and selected in 2Bi, Bii by qPCR: CABG-NDM, CABG-DM, MVR, AVR, peripheral blood
*(n = 12 samples per group, up to 4 time points; up to 192 samples total)*

iv. Proteomics analysis: pre-operative blood and matched transcoronary blood
*(n = 3-5 samples per group, as for Bi)*
Subsequent time-points if funding obtained.

v. Metabolomics analysis (Marc-Emmanuel Dumas - Imperial College London): pre-operative blood and matched transcoronary blood
*(n = 3-5 samples per group, as for Bi)*
Subsequent time-points if funding obtained.

**C. Peri-operative blood plasma extracellular vesicles (EVs) – exosomes and microparticles (MPs)**

i. qPCR analyses of the ncRNAs emerging from the analyses in the whole plasma (part 2B) at the level of exosomes and microparticle fractions of the plasma (initial focus on transcoronary and arterial line blood followed by time course in peripheral blood): CABG-NDM, CABG-DM, MVR, AVR
*(Up to 216 samples total**)*

ii. Proteomics analysis: EVs from samples as in 2Ci.

iii. Metabolomics analysis: EVs from samples as in 2Ci.

D. **Pericardial fluid**

i. miRNA qPCR arrays on whole PF: CABG-NDM, CABG-DM, MVR, AVR (matched with transcoronary bloods and LV)
*(n = 3-5 per group*. One group has 3 samples, two groups have 4 samples each, and 1 group has 5 samples.)*

ii. qPCR validation of selected findings from LV biopsy sequencing and qPCR array(s): CABG-NDM, CABG-DM, MVR, AVR
*(12 samples per group; 48 samples total)*

iii. Proteomics analysis: CABG-NDM, CABG-DM, MVR, AVR
*(n = 12 per group; 48 samples total)*

iv. Metabolomics analysis: CABG-NDM, CABG-DM, MVR, AVR
*(n = 12 per group; 48 samples total)*

E. **Pericardial fluid extracellular vesicles (EVs) – exosomes and microparticles (MPs)**

i. miRNA qPCR arrays: CABG-NDM, CABG-DM, MVR, AVR with matched transcoronary bloods
*(n = 3-5 per group*. One group has 3 samples, two groups have 4 samples each, and 1 group has 5 samples.)*

ii. qPCR validation of selected findings from LV biopsy sequencing and qPCR array(s): CABG-NDM, CABG-DM, MVR, AVR
*(12 samples per group; 48 samples total)*

iii. Proteomics analysis: EVs from samples as in 2Diii.

iv. Metabolomics analysis: EVs from samples as in 2Div.

* original design was 6 per group but some samples unavailable.

** original design was 216 samples (4 groups; 6 patients per group with 4 samples, 6 patients per group with 5 samples) but recent experience suggests some will be unavailable.
