## Supplementary file 2 for "Human left ventricle circRNA-miRNA-mRNA network analyses reveals a novel proangiogenic role for circNPHP1 under ischemic conditions"

### Supplementary File 2: Resources Table

***Antibodies***

| Target antigen | Vendor or Source | Catalog # | Working concentration | Lot # (preferred but not required) | Persistent ID / URL |
|---|---|---|---|---|---|
| VEGF-A (WB) | Abcam | ab46154 | 1:2000 | 1051635-1 | https://www.abcam.com/products/primary-antibodies/vegfa-antibody-ab46154.html |
| BCL2 (WB) | Cell Signalling Technology | 3498S | 1:1000 | 6 | https://www.cellsignal.com/products/primary-antibodies/bcl-2-d17c4-rabbit-mab/3498 |
| Lamin B1 (WB) | Cell Signalling Technology | 12586S | 1:1000 | 2 | https://www.cellsignal.com/products/primary-antibodies/lamin-b1-d4q4z-rabbit-mab/12586 |
| Mouse-anti-rabbit IgG-HRP (WB) | Santa Cruz | sc-2357 | 1:2000 | H2218 | https://www.scbt.com/p/mouse-anti-rabbit-igg-hrp |
| Phalloidin-FITC (Matrigel assay) | ThermoFisher Scientific | A12379 | 1:2500 | | https:// www.thermofisher.com/order/catalog/product/A12379 |

***Cultured Cells***

| Name | Vendor or Source | Persistent ID / URL |
|---|---|---|
| Human Umbilical Vein Endothelial Cells 2 (HUVEC 2) (pooled donors, cryopreserved vials) | Promocell | Catalogue no: C-12208<br>https://promocell.com/product/human-umbilical-vein-endothelial-cells-2-huvec-2/<br><br>Lot numbers: 447Z004, 450Z015, 466Z022, 503Z026 |
| Human Cardiac Microvascular Endothelial Cells (HCMEC) (single donors, cryopreserved vials) | Promocell | Catalogue no: C-12285<br>https://promocell.com/product/human-cardiac-microvascular-endothelial-cells-hcmec/<br><br>Lot numbers: 440Z021.5, 440Z021.4, 446Z001.1, 492Z009.4 |
| Human Cardiac Fibroblasts (HCF), unknown if pooled or single donor cryopreserved vials | Promocell | Catalogue no: C-12375<br>https://promocell.com/product/human-cardiac-fibroblasts-hcf/<br><br>Lot number: 491Z026.1 |
| Human ventricular cardiomyocyte (AC-16 cells) | Donated by collaborator (Dr Rajesh Katare University of Otago) | https://www.sigmaaldrich.com/GB/en/product/mm/scc109 |

***Data Availability***

| Description | Source / Repository | Persistent ID / URL |
|---|---|---|
| LV biopsies whole transcriptome RNA sequencing | GEO database | In progress |
| LV biopsies small RNA sequencing | GEO database | In progress |
| Single cell sequencing data of Human Heart Atlas | The European Nucleotide Archive (ENA) database<br><br>Processed data exploration from Heart Cell Atlas v2 website | ERP123138<br><br>https://www.heartcellatlas.org |

| | | |
|---|---|---|
| Single cell sequencing data of normal and diabetic human hearts | GEO database | GSE183852 |
| RNA sequencing data on circular RNAs (HUAECs) | GEO database | GSE100242 |
| Expression profiling of miRNAs (HCAECs) by RT-PCR | GEO database | GSE53315 |
| Transcriptomic profiling by RNA-seq (HCAECs) | GEO database | GSE134489 |

***List of oligos***

| Description | Sequence | Vendor | Persistent ID / URL |
|---|---|---|---|
| Control siRNA | ON-TARGETplus Non-targeting Pool | Horizon Discovery Catalog# D-001810-10-05 | https://horizondiscovery.com/en/gene-modulation/knockdown/sirna/products/on-targetplus-sirna-reagents |
| CircNPHP1 siRNA | Sense: 5'AGAAGGCACUGAUAAACACUGUUU 3' Antisense: 5'ACAGUGUUUAUCAGUGCCUUCUUU 3' | Horizon Discovery (Custom made) | - |
| Human has-miR-221-3p (microRNA Mimic) | See URL | Horizon Discovery Catalog# C-300578-05-0010 | https://horizondiscovery.com/en/gene-modulation/knockdown/mirna/products/miridian-microrna-mimic?nodeid=mirnaprecursor-mi0000298 |
| Mimic Negative Control #1 | See URL | Horizon Discovery Catalog# CN-001000-01-05 | https://horizondiscovery.com/en/gene-modulation/knockdown/controls/products/miridian-microrna-mimic-negative-control-1 |
| Anti-miR™ miRNA Inhibitor miR-221-3p | See URL | ThermoFisher Scientific Catalog# AM17000 | https:// www.thermofisher.com/order/catalog/product/AM17000 |
| Anti-miR™ miRNA inhibitor Negative Control #1 | See URL | ThermoFisher Scientific Catalog# AM17010 | https:// www.thermofisher.com/order/catalog/product/AM17010 |
| cel-miR-39 (microRNA (cel-miR-39) Spike-In Kit) | See URL | Norgen Biotek Catalog# SKU 59000 | https://norgenbiotek.com/product/microrna-cel-mir-39-spike-kit |
| Control ASO Probe: | 5'TGCGTAACGAACGACGAATCGTCGCAGATC-3'[Biotin-TEG] | Sigma-Aldrich (Custom made) | - |
| circNPHP1 ASO Probe1: | 5'CTACAGAAGGCACTGATAAACACTGTTGAT-3'[Biotin-TEG] | Sigma-Aldrich (Custom made) | - |
| circNPHP1 ASO Probe2**:** | 5'AGAAGGCACTATAAACACTGT-3'[Biotin-TEG] | Sigma-Aldrich (Custom made) | - |

***List of qRT-PCR primers***

| Description | Sequence | Vendor |
|---|---|---|
| Linear NPHP1 | Forward 5’ CTGCCACTGTCACACTGCATTC 3’<br>Reverse 5’ GTTTACCATGACTGCGTGCTCC 3’ | Sigma-Aldrich |
| CircNPHP1 | Forward 5’ TCTTACAACCAGAGCTCATGCC 3’<br>Reverse 5’ AAGCTGTGAGAGCGTGGAA | Sigma-Aldrich |
| VEGFA | Forward 5’ AGAGCAAGACAAGAAAATCC 3’<br>Reverse 5’ TACAAACAAATGCTTTCTCC 3’ | Sigma-Aldrich |
| BCL2 | Forward 5’ ACTGGAGAGTGCTGAAGATTG 3’<br>Reverse 5’ AGTCTACTTCCTCTGTGATGTTG 3’ | Sigma-Aldrich |
| BCL2L11 (BIM) | Forward 5’ GGCCCCTACCTCCCTACA 3’<br>Reverse 5’ GGGGTTTGTGTTGATTTGTCA 3’ | Sigma-Aldrich |
| 18s | Forward 5’ CCCAGTAAGTGCGGGTCAT 3’<br>Reverse 5’ CCGAGGGCCTCACTAAACC 3’ | Sigma-Aldrich |
| GAPDH | Forward 5’ GACTCATGACCACAGTCCATGC 3’<br>Reverse 5’ AGAGGCAGGGATGATGTTCTG 3’ | Sigma-Aldrich |
| cel-miR-39 | Forward 5’ TTGCAGCTCTCATAGAAGGAACCG3′<br>Reverse 5′-GTTTCAGCCGAGACTAGACTTTGAGC3′ | Sigma-Aldrich |
| CircMCU | Forward 5’CTGTTCACGCAGGGGAAACT 3’<br>Reverse 5’ AGCAGCTAAGATGTCACTGGC 3’ | Sigma-Aldrich |
| CircSTX17 | Forward 5’ ATGCTGCAGAATCGTGGGAA 3’<br>Reverse 5’ TCTGAGAACTAGCTTCAGCTTCA 3’ | Sigma-Aldrich |
| CircMGA | Forward 5’ TGTAAGCCCTGGGAGTACCT 3’<br>Reverse 5’ TCTGGTCTAACGGTGAGGCT3’ | Sigma-Aldrich |
| hsa-miR-221-3p | hsa-miR-221-3p miRCURY LNA miRNA PCR Assay (predesigned) GeneGlobe ID - YP00204532 | Qiagen<br>Catalog# 339306 |
| hsa-miR-222-3p | hsa-miR-222-3p miRCURY LNA miRNA PCR Assay (predesigned) GeneGlobe ID - YP00204551 | Qiagen<br>Catalog# 339306 |
| hsa-miR-299-3p | hsa-miR-299-3p miRCURY LNA miRNA PCR Assay (predesigned) GeneGlobe ID - YP00204702 | Qiagen<br>Catalog# 339306 |
| hsa-miR-139-3p | hsa-miR-139-3p miRCURY LNA miRNA PCR Assay (predesigned) GeneGlobe ID - YP00205661 | Qiagen<br>Catalog# 339306 |
| hsa-miR-141-3p | hsa-miR-141-3p miRCURY LNA miRNA PCR Assay (predesigned) GeneGlobe ID - YP00204504 | Qiagen<br>Catalog# 339306 |
| U6 | U6 snRNA (v2) miRCURY LNA miRNA PCR Assay<br>(predesigned) GeneGlobe ID - YP00204532 | Qiagen<br>Catalog# 339306 |

***Others***

| Description | Source / Repository | Persistent ID / URL |
|---|---|---|
| Endothelial cell growth medium 2 | Promocell | https://promocell.com/product/endothelial-cell-growth-medium-2/ |
| Endothelial cell growth medium MV | Promocell | https://promocell.com/product/endothelial-cell-growth-medium-mv/ |
| Fibroblast Growth Medium 3 | Promocell | https://promocell.com/product/fibroblast-growth-medium-3/ |
| DMEM-F12 | ThermoFisher Scientific | https://www.thermofisher.com/order/catalog/product/10565018 |
| Lipofectamine™ 2000 | ThermoFisher Scientific | https://www.thermofisher.com/order/catalog/product/11668027 |
| BrdU Cell Proliferation ELISA kit | Abcam | https://www.abcam.com/products/elisa-kits/brdu-cell-proliferation-elisa-kit-colorimetric-ab126556.html |
| Growth Factor Reduced Matrigel | Corning | https://ecatalog.corning.com/life-sciences/b2b/DK/en/Surfaces/Extracellular-Matrices-ECMs/Corning%C2%AE-Matrigel%C2%AE-Matrix/p/356231 |
| RealTime-Glo™ Annexin V Apoptosis and Necrosis Assay kit | Promega | https://www.promega.co.uk/products/cell-health-assays/apoptosis-assays/realtime-glo-annexin-v-apoptosis-assay/?catNum=JA1011 |
| Streptavidin Dynabeads | New England Biolabs | https://www.neb.com/en-gb/products/s1420-streptavidin-magnetic-beads |
| miRNeasy mini kit | Qiagen | https://www.qiagen.com/us/products/discovery-and-translational-research/dna-rna-purification/rna-purification/mirna/mirneasy-kits?catno=217004 |
| QIAzol reagent | Qiagen | https://www.qiagen.com/us/products/discovery-and-translational-research/lab-essentials/buffers-reagents/qiazol-lysis-reagent |
| miRNeasy Serum/Plasma Kit | Qiagen | https://www.qiagen.com/us/products/discovery-and-translational-research/dna-rna-purification/rna-purification/mirna/mirneasy-serumplasma-kit |
| PrimeScript RT-PCR kit | TakaraBio | https://www.takarabio.com/products/real-time-pcr/reverse-transcription-prior-to-qpcr/convenient-master-mix-for-real-time-pcr |
| TB Green Premix Ex Taq II Kit | TakaraBio | https://www.takarabio.com/products/real-time-pcr/real-time-pcr-kits/dye-based-qpcr-mixes/tb-green-premix-ex-taq-ii-(tli-rnase-h-plus)?catalog=RR82LR |
| miRCURY LNA RT Kit | Qiagen | https://www.qiagen.com/us/products/discovery-and-translational-research/pcr-qpcr-dpcr/qpcr-assays-and-instruments/mirna-qpcr-assay-and-panels/mircury-lna-rt-kit |
| miRCURY LNA SYBR Green PCR Kit | Qiagen | https://www.qiagen.com/us/products/discovery-and-translational-research/pcr-qpcr-dpcr/qpcr-assays-and-instruments/mirna-qpcr-assay-and-panels/mircury-lna-sybr-green-pcr-kits |
| Immobilon Crescendo Western HRP | Merck | https://www.merckmillipore.com/GB/en/product/Immobilon-Crescendo-Western-HRP-substrate-100-mL,MM_NF-WBLUR0100 |

| | | |
|---|---|---|
| Protease inhibitor cocktail | Merck | https://www.sigmaaldrich.com/GB/en/product/roche/11836153001?utm_source=google&utm_medium=cpc&utm_campaign=15001183107&utm_content=127306761543&gclid=CjwKCAjwwr6wBhBcEiwAfMEQsyfkKcrb8eE7ZpKtmtuk13a6ja4dLQZOrlOxIenI4CT6Y6NEL0S5txoC7ZQQAvD_BwE |
| RIPA buffer | Sigma-Aldrich | https://www.sigmaaldrich.com/GB/en/product/sigma/r0278 |
| *mir*Vana kit (AM1560) | ThermoFisher Scientific | https://www.thermofisher.com/order/catalog/product/AM1560?gclid=CjwKCAjwwr6wBhBcEiwAfMEQsw4Pcovfnp_wS2LJKczpGy-I84MvofdhpFVR95-G-g11aQJ4QnHF0hoC2KYQAvD_BwE&ef_id=CjwKCAjwwr6wBhBcEiwAfMEQsw4Pcovfnp_wS2LJKczpGy-I84MvofdhpFVR95-G-g11aQJ4QnHF0hoC2KYQAvD_BwE:G:s&s_kwcid=AL!3652!3!606515189508!!!g!!!16893189581!143415109412&cid=bid_sap_rst_r01_co_cp0000_pjt0000_bid00000_0se_gaw_dy_awa_con&gad_source=1 |
| DNA-*free* DNA removal kit | Fisher Scientific | https://www.fishersci.com/shop/products/ambion-dna-i-free-i-dna-removal-kit-1/AM1906 |
| NEBNext rRNA depletion kit | New England Biolabs | https://www.neb.com/en-gb/products/e6350-nebnext-rrna-depletion-kit-human-mouse-rat-with-sample-purification-beads |
| NEBNext Ultra II Directional RNA Library Prep Kit for Illumina | New England Biolabs | https://www.neb.com/en-gb/products/e7760-nebnext-ultra-ii-directional-rna-library-prep-kit-for-illumina |
