## Supplementary table 1 for "Human left ventricle circRNA-miRNA-mRNA network analyses reveals a novel proangiogenic role for circNPHP1 under ischemic conditions"

| Patients Characteristics | | | |
|---|---|---|---|
| | IHD (CABG) [n=12] | IHD+T2DM (CABGD) [n=11] | Non-IHD (MVR) [n=12] |
| **Age (years)** | 41-76 | 54-77 | 46-76 |
| **Sex** | 10 M ; 2 F | 9 M ; 2 F | 7 M ; 5 F |
| **Weight (Kg)** | 93.2 (± 30) | 86.1 (±30) | 82.3 (±20) |
| **Diabetes (%)** | 0 | 100 | 0 |
| **Heart Failure (NYHA %)** | I (25) II (75) III (0)VI (0) | I (18) II (36) III (46) VI (0) | I (33) II (42) III (25) VI (0) |
| **LV function (%)** | 1 (92) 2(8) 3(0) | 1 (64) 2(36) 3(0) | 1 (92) 2(8) 3(0) |
| **Previous MI (%)** | 25 | 36 | 0 |
| **Smoker (%)** | 0 / Ex-smoker (33) | 0 / Ex-smoker (54) | 0 / Ex-smoker (33) |
| **CVD family history (%)** | 67 | 82 | 33 |
| **Hypertension (%)** | 84 | 100 | 58 |
| **Hypercholesterolemia (%)** | 84 | 82 | 67 |
| **Hyperthyroidism (%)** | 16 | 0 | 0 |
| **Preoperative medications (%)** | | | |
| ***Aspirin*** | 83 | 90 | 17 |
| ***Clopidogrel*** | 33 | 54 | 0 |
| ***Beta blockers*** | 83 | 82 | 33 |
| ***Oral Nitrates*** | 50 | 9 | 8 |
| ***Statin*** | 92 | 82 | 58 |
| ***ACE inhibitors*** | 42 | 64 | 42 |
| ***Angiotensin blockers*** | 0 | 9 | 8 |
| ***Diuretics*** | 1 | 36 | 33 |
| ***Oral anti-diabetics*** | 0 | 54 | 0 |
| ***Insulin*** | 0 | 27 | 0 |

***Legend***

***Sex***

| | |
|---|---|
| **M** | **Male** |
| **F** | **Female** |

***LV function***

| | |
|---|---|
| **1** | **Good (>50%)** |
| **2** | **Moderate (30-50%)** |
| **3** | **Poor (<30%)** |
